# Sec7 regulatory domains scaffold autoinhibited and active conformations

**DOI:** 10.1101/2023.11.22.568272

**Authors:** Bryce A. Brownfield, Brian C. Richardson, Steve L. Halaby, J. Christopher Fromme

## Abstract

The late stages of Golgi maturation involve a series of sequential trafficking events in which cargo-laden vesicles are produced and targeted to multiple distinct subcellular destinations. Each of these vesicle biogenesis events requires activation of an Arf GTPase by the Sec7/BIG guanine nucleotide exchange factor (GEF). Sec7 localization and activity is regulated by autoinhibition, positive feedback, and interaction with other GTPases. Although these mechanisms have been characterized biochemically, we lack a clear picture of how GEF localization and activity is modulated by these signals. Here we report the cryoEM structure of full-length Sec7 in its autoinhibited form, revealing the architecture of its multiple regulatory domains. We use functional experiments to determine the basis for autoinhibition and use structural predictions to produce a model for an active conformation of the GEF that is supported empirically. This study therefore elucidates the conformational transition that Sec7 undergoes to become active on the organelle membrane surface.

## Introduction

The Golgi apparatus is the central organelle of the secretory pathway in eukaryotic cells, connecting synthesis of proteins and lipids at the endoplasmic reticulum (ER) with the plasma membrane (PM) and endolysosomal system. Approximately 30% of the eukaryotic proteome requires the Golgi for post-translational modifications and sorting to function at specific locations. Vectoral flow of cargo through the Golgi requires timely and accurate packaging into transport vesicles and modification of the membrane lipid constituents. Numerous pathway-specific factors are recruited to and activated on the Golgi membrane at a precise time and location. The final stage of the Golgi, the *trans*-Golgi network (TGN), connects the Golgi to the PM and endolysosomal system. Significant changes to the membrane itself occur at the TGN, and multiple trafficking pathways originate from this dynamic compartment.

Membrane trafficking is regulated by Ras-related small GTPases of the Arf and Rab families (1, 2). These GTPases act as molecular switches by transitioning between soluble GDP-bound and membrane-anchored GTP-bound conformations to recruit and activate effectors on organelle and vesicle membranes. The intrinsic conversion between these two states is essentially negligible, occurring less frequently than once every 10^5^ seconds (3, 4). GEFs facilitate GDP to GTP exchange by displacing bound nucleotide, driven by a higher cytosolic GTP concentration, and ‘GTPase activating proteins’ (GAPs) inactivate GTPases by stimulating GTP hydrolysis to convert bound GTP to GDP. Coordination between GEFs, GAPs, GTPases, and effectors at the Golgi gives rise to an ordered progression through a series of trafficking pathways referred to as Golgi maturation (5).

At the Golgi, Arf1 and its close paralogs (Arf1-5 in mammals, Arf1-2 in yeast) are essential for vesicle biogenesis by recruiting cargo adaptors, membrane modifying enzymes, and coat proteins. Arf GTPases can also directly induce curvature by inserting an amphipathic helix into the outer leaflet of the membrane (Figure 1a) (2, 6, 7). In the budding yeast (*Saccharomyces cerevisiae*) model system, three “large” Arf-GEFs activate Arf1 at the early-, medial-, and late-Golgi/TGN: Gea1, Gea2, and Sec7, respectively (8–10). In mammals, GBF1 is homologous to Gea1/2 and the BIG/ARFGEF proteins are homologous to Sec7 (11, 12).

**Figure 1.**
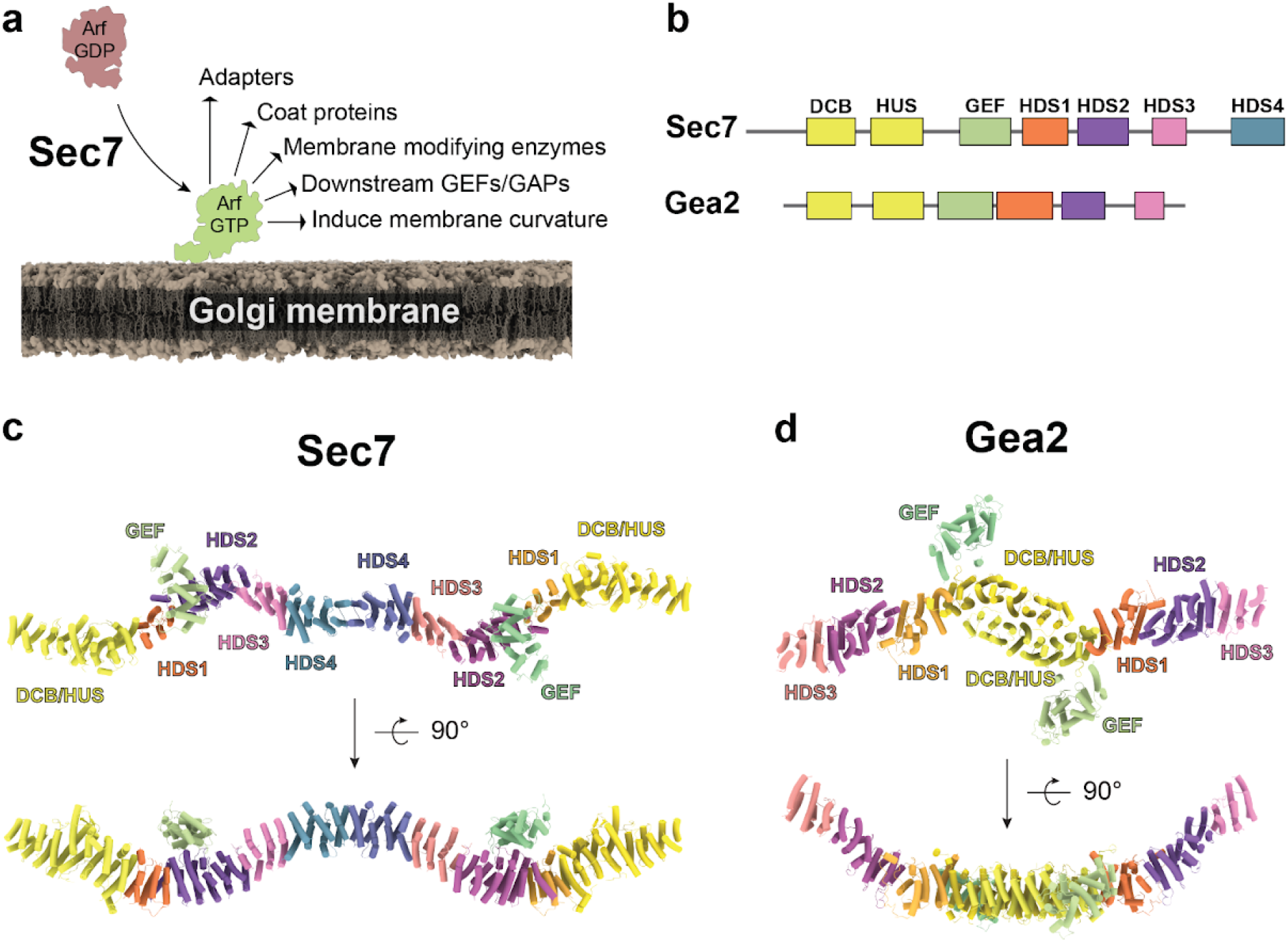
CryoEM structure of Sec7. **a,** Sec7 activates Arf1 on the TGN membrane to trigger many vesicle trafficking functions. **b**, Domain architecture of Sec7 and Gea2. **c**, CryoEM model for the full Sec7 dimer, colored as in (b). **d**, CryoEM model of the Gea2 “open” dimer PDB: 7UTH (19), colored as in (b). Note the difference in dimerization via the HDS4 domain (Sec7, c) and DCB-HUS domain (Gea2, d).

Sec7 is regulated by autoinhibition and a network of GTPase interactions including positive feedback (13, 14). Sec7 possesses six identified regulatory domains (15) that mediate its localization and regulation. Although these domains are conserved across Sec7 homologs in all eukaryotes, they are not found in other proteins and therefore it has not been possible to infer their function from homology alone. Previous work from our laboratory indicated the Sec7 HDS1-4 domains are autoinhibitory, the DCB-HUS domain is important for activating Arf1 on the membrane surface, the ‘HUS-box’ is important for allosteric stimulation of Sec7, and the HDS1 domain is important for positive feedback (13–17). However, how these different regulatory behaviors are integrated by Sec7 is not known. To understand how the regulatory domains of Sec7 cooperate to regulate its localization and activity, structural information is necessary.

Here we report the structure of full-length Sec7 determined by cryogenic electron microscopy (cryoEM) to 3.7 Å average resolution. The structure reveals the overall architecture of the regulatory and GEF domains within the context of the Sec7 homodimer. The arrangement of the Sec7 dimer is strikingly different than that of the related Arf-GEF Gea2. Whereas Gea2 dimerizes via its N-terminal DCB-HUS domain, Sec7 dimerizes via its C-terminal HDS4 domain. The structure also indicates that the primary mechanism for autoinhibition is an interaction between the GEF and HDS2 domains that occludes the catalytic surface of the GEF domain. An alternative conformation of the GEF domain is predicted by AlphaFold (18) and resembles the positioning of the GEF domain in Gea2 (19). As Gea2 is not autoinhibited, we infer that the predicted structural conformation represents an active state of Sec7. We also performed extensive *in vitro* and *in vivo* functional experiments to test and validate both the cryoEM and predicted structural models. Our findings enable us to provide a structural explanation for how the Sec7 regulatory domains function in both the autoinhibited and active states.

## Results

### CryoEM structure of *T. terrestris* Sec7

To determine how Arf activation at the TGN is regulated by the Arf-GEF regulatory domains (Figure 1a,b), we sought the structure of an intact Sec7/BIG homolog. We therefore purified full-length Sec7 from the thermophilic yeast *Thielavia terrestris* using overexpression in *Pichia pastoris*, verified that it possessed *in vitro* GEF activity towards Arf1, and determined its structure by single-particle cryoEM (Figure 1c, Tables 1 and 2, SI Appendix, Fig. S1, and SI Appendix, Video S1).

**Table 1.**
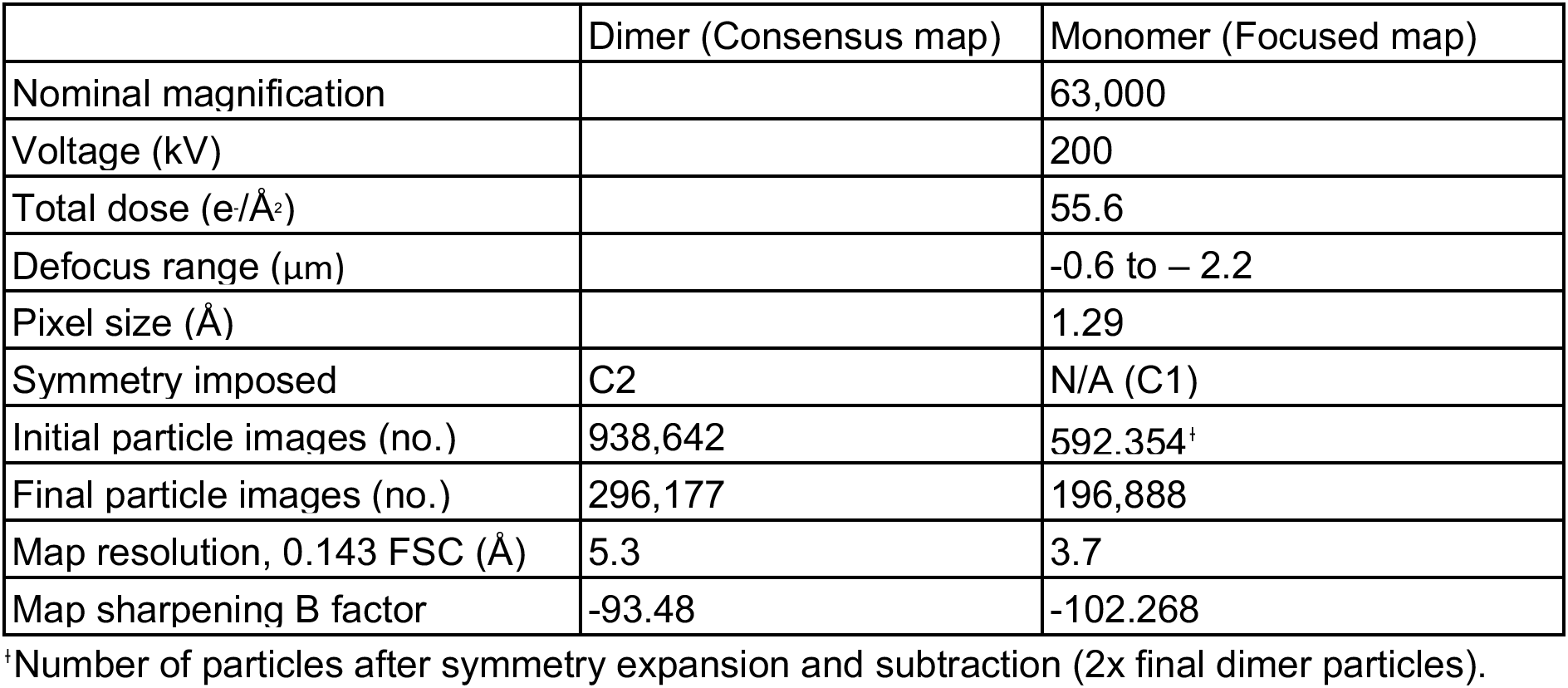
CryoEM data collection, processing, and statistics.

**Table 2.**
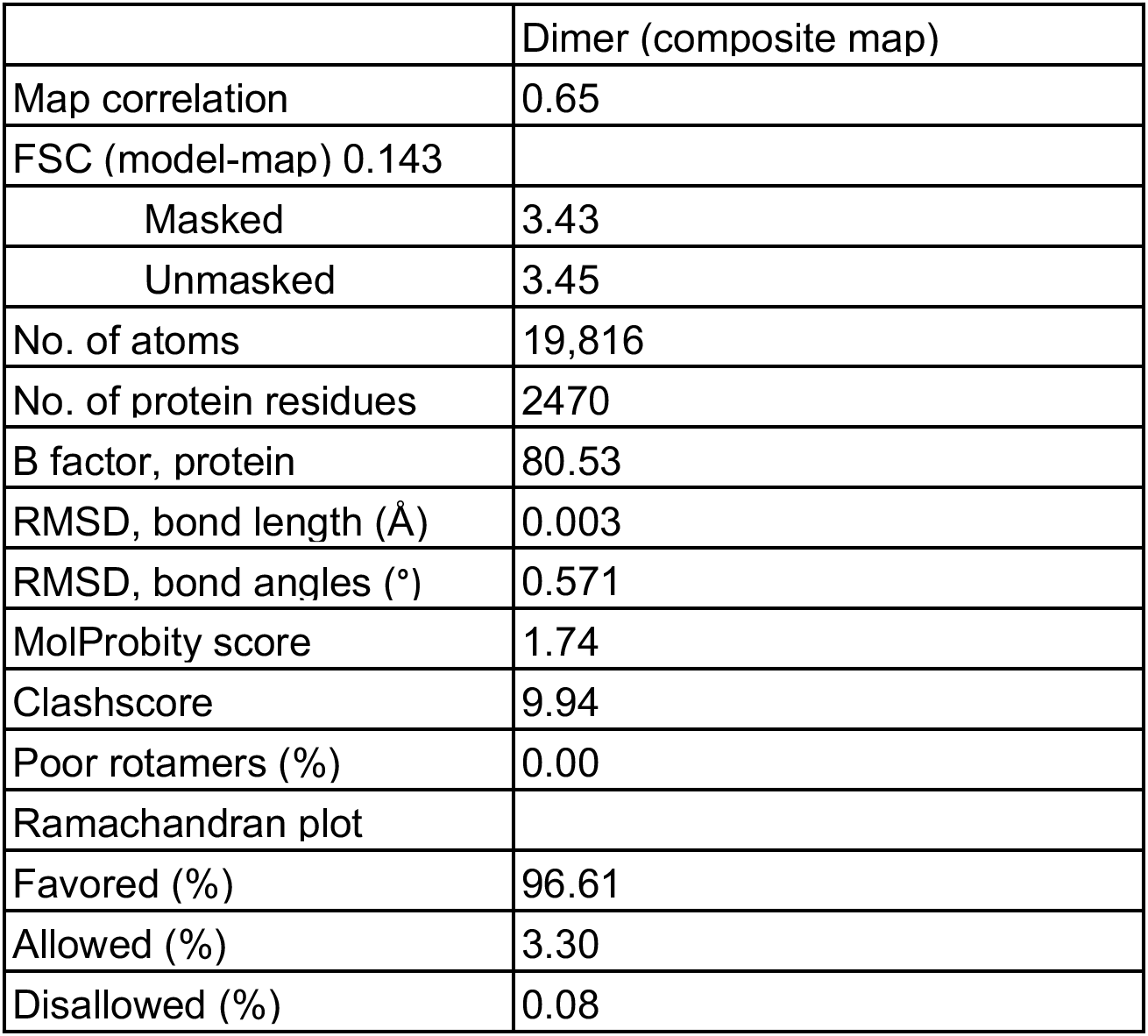
Model statistics.

|  | Dimer (composite map) |
| --- | --- |
| Map correlation | 0.65 |
| FSC (model-map) 0.143 |  |
| Masked | 3.43 |
| Unmasked | 3.45 |
| No. of atoms | 19,816 |
| No. of protein residues | 2470 |
| B factor, protein | 80.53 |
| RMSD, bond length (Å) | 0.003 |
| RMSD, bond angles (°) | 0.571 |
| MolProbity score | 1.74 |
| Clashscore | 9.94 |
| Poor rotamers (%) | 0.00 |
| Ramachandran plot |  |
| Favored (%) | 96.61 |
| Allowed (%) | 3.30 |
| Disallowed (%) | 0.08 |

Exploiting the C2 symmetry of the Sec7 dimer allowed us to generate a preliminary 5.3 Å resolution density map (SI Appendix, Fig. S1d), however significant flexibility across the thin ∼40 nm molecule limited the resolution, especially at the distal ends of the dimer. Heterogeneous reconstruction analysis by cryoDRGN (20) suggested each end of the molecule can flex as much as 28° relative to the central dimerization domain (SI Appendix, Fig. S2). This flexibility is apparently continuous, as intermediate states were poorly reconstructed and standard real space 3D classification was unable to isolate discrete states.

Subsequent symmetry expansion, particle subtraction, and 3D classification of individual monomers enabled us to generate a significantly improved reconstruction of the monomer with an average 0.143 FSC of 3.7Å (SI Appendix, Fig. S3). Focused refinements of finer sub- volumes did not further improve resolution, so the monomer map was used to build and refine an atomic model of a Sec7 monomer, which was then combined and refined into a composite dimer map to produce the full model of a Sec7 homodimer (Figure 1c, SI Appendix, Fig. S1d).

The architecture of the Sec7 dimer is a long continuous alpha-solenoid consisting of tandem pairs of α-helices forming an elongated ladder structure we refer to as the backbone (Figure 1c). The six previously described regulatory domains of Sec7 (DCB, HUS, HDS1-4) comprise the backbone. The catalytic GEF domain, which lies in between the HUS and HDS1 domains in the primary sequence, is extruded from the backbone such that the HUS and HDS1 domains directly interact with each other. The GEF domain is bound to the surface of the HDS2 domain. Homo-dimerization is mediated by the HDS4 domain, consistent with conclusions from previous studies (16, 21) but distinct from the dimerization mode of the related Arf-GEF Gea2 (19, 22). As a result of dimerization Sec7 adopts a flattened ‘W’ architecture when viewed from the side, and a sinuous partial corkscrew when viewed from above (Figure 1c,d).

### Autoinhibition of Sec7 via the HDS2 domain

The cryoEM structure reveals the HDS2 domain interacts with the catalytic surface of the GEF domain in a manner that is mutually exclusive with Arf1 binding. When we superimposed the crystal structure of the Gea2 GEF domain bound to nucleotide-free Arf1 (23) onto the GEF domain in the Sec7 cryoEM structure, we observed a significant steric clash between Arf1 and the HDS2 domain (Figure 2a). This indicates that the observed position of the GEF domain bound to the HDS2 domain prevents Arf1 activation, providing a clear mechanism for autoinhibition. We previously reported that truncations of the C-terminal HDS2-HDS4 domains resulted in a construct with higher *in vitro* GEF activity (8, 13, 14), consistent with the observed HDS2-GEF domain interaction serving to mediate autoinhibition.

**Figure 2.**
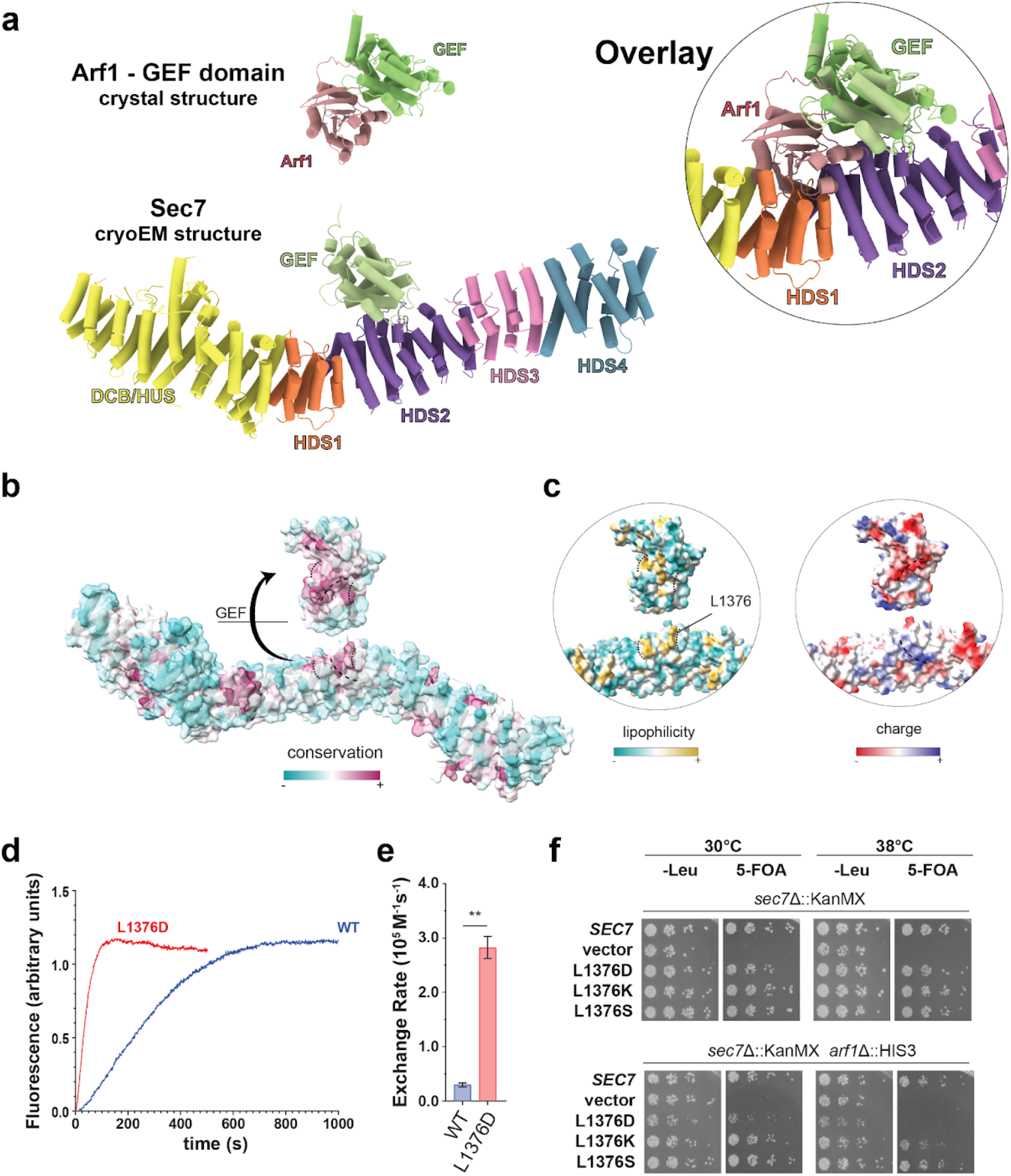
The GEF:HDS2 interaction interface is autoinhibitory and physiologically important. **a,** Crystal structure of the Gea2 GEF domain bound to Arf1(23) superimposed onto the Sec7 cryoEM structure (inset) illustrating how the HDS2 domain prevents Arf1 from interacting with the GEF domain. **b**, Sec7 model surface colored by Consurf (29) analysis showing the HDS2:GEF interface is well conserved. **c**, Surface maps of the model as shown in (b) colored by lipophilicity (left) and electrostatic potential (right). The site of the mutation is shown and labeled according to *S. cerevisiae* residue numbering (the *T. terrestris* equivalent residue is L1177). **d,** Sec7 GEF activity assay results showing a representative trace of Tryptophan fluorescence after addition of GTP (see Methods) to monitor Arf activation kinetics *in vitro*. **e**, Average quantified rate constant of three replicate GEF assays. **f**, Complementation test (plasmid shuffling assay) indicating L1376D Sec7 is able to support normal growth in the *sec7*Δ strain, but not the sensitized *sec7*Δ*arf1*Δ strain.

The arrangement of the GEF domain in Sec7 is quite different than that observed in the cryoEM structure of Gea2 (19) (Figure 1c,d), in which the GEF domain is positioned adjacent to the DCB-HUS domain in a manner that does not interfere with Arf1 binding. Unlike Sec7, Gea1 and Gea2 are not autoinhibited (8) and do not appear to be regulated by positive feedback (13). We compared the sequences of Sec7 and Gea2 in the region of the HDS2 domain contacting the GEF domain and found these sequences to be well conserved across Sec7 homologs but significantly less well conserved in Gea homologs (Figure 2b, SI Appendix, Fig. S4a). On the GEF domain-binding surface of HDS2, two loops of predominantly hydrophobic conserved residues flank a weakly basic groove to form a surface that is complementary to similarly well conserved regions of the GEF domain (Figure 2b,c, SI Appendix, Fig. S4b). We note this interaction is different from the autoinhibitory interaction of the cytohesin Arf-GEF Grp1 in which a basic patch in the PH domain C-terminus blocks the catalytic surface of the Grp1 GEF domain (24) (SI Appendix, Fig. S4c).

To test the importance of the HDS2-GEF domain interaction for the autoinhibitory behavior of Sec7, we engineered a mutation in *S. cerevisiae* Sec7 designed to disrupt the exposed hydrophobic surface of the HDS2 domain. We purified this L1376D mutant *S. cerevisiae* Sec7 protein, along with the wild-type, and tested its ability to activate Arf1 on a membrane surface using an established *in vitro* GEF activity assay (13, 25). A change in the native tryptophan fluorescence of Arf1 can be used to monitor activation kinetics of Arf family proteins (24, 26–28). Using myristoylated-Arf1 and synthetic liposomes with a lipid mixture mimicking that of the TGN, we observed that the purified L1376D Sec7 mutant construct exhibited significantly higher GEF activity than wild-type Sec7 (Figure 2d,e, SI Appendix, Fig. S4f). This result supports our interpretations that the HDS2-GEF domain interface mediates autoinhibition of Sec7, and that the cryoEM structure represents the autoinhibited conformation of Sec7.

To determine the *in vivo* significance of this autoinhibitory interaction, we tested the ability of Sec7 constructs harboring mutations in L1376 to complement the loss of *SEC7* gene function in *S. cerevisiae* cells. We determined that in a sensitized *arf1*Δ background, in which only 10% of the total Arf1/2 protein remains (30), the L1376D mutation did not reduce expression but reduced viability at 30 °C and was lethal at 38 °C (Figure 2f, SI Appendix, Fig. S4d). In contrast, the L1376K and L1376S mutations did not have significant impacts on growth. Wild-type Sec7 localizes to late-Golgi/TGN compartments which appear as puncta distributed in the cytosol(9, 13, 31). We visualized the localization of the L1376D mutant Sec7 protein in live cells by fluorescence microscopy and observed that the mutation did not disrupt its localization to punctate structures (SI Appendix, Fig. S4e). Interestingly, the morphology of the Sec7 labeled Golgi compartments in the mutant cells appeared significantly more variable in size and intensity, consistent with an impact on Golgi function. Taken together, although the L1376D mutation was tolerated in the wild-type ARF1 strain, these results indicate that disruption of the autoinhibitory HDS2-GEF domain interface is deleterious *in vivo*.

### AlphaFold predicts an active conformation of Sec7

We compared the AlphaFold (18) prediction of *S. cerevisiae* Sec7 to our experimental cryoEM model and found a striking difference in the position of the GEF domain between the two models. Rather than binding to the HDS2 domain as we observed in the experimental structure, AlphaFold predicted the GEF domain to occupy a position that closely resembles the “open” conformation observed in the cryoEM structure of Gea2 (19) (Figure 3a). Importantly, in the predicted Sec7 structure the GEF domain is available to bind Arf1 without steric hindrance (SI Appendix, Fig. S5a). Furthermore, linker regions between the GEF and backbone domains make extensive contact with the “HUS box” (SI Appendix, Fig. S5b), a conserved motif near the C-terminal end of the HUS domain implicated in allosteric stimulation of Sec7 GEF activity (15, 17). We therefore considered the AlphaFold prediction to represent a plausible model for the active conformation of the Sec7 GEF domain.

**Figure 3.**
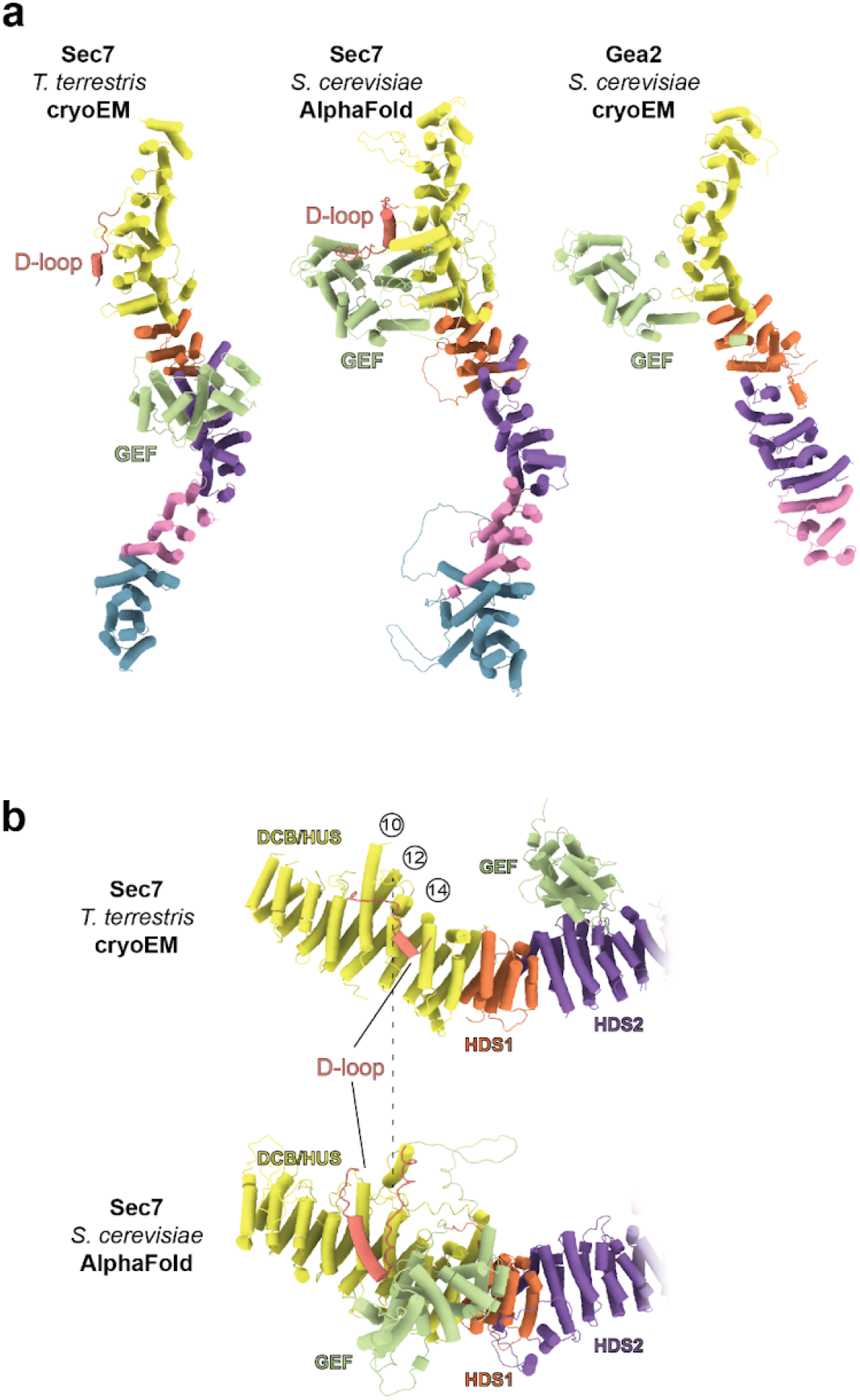
AlphaFold predicts an active conformation of Sec7. **a**, Side by side comparison of the *T. terrestris* Sec7 cryoEM structure, the AlphaFold predicted *S. cerevisiae* Sec7 model, and the cryoEM structure of *S. cerevisiae* Gea2 in the “open” conformation (PDB: 7UTH). **b**, The “D-loop” (colored salmon) binds to a surface between helices 12 and 14 of the DCB-HUS domain in the Sec7 cryoEM structure, but is displaced by the GEF domain and instead bound to the surface of helices 10 and 12 in the AlphaFold prediction.

In this conformation the GEF domain has displaced a partially ordered loop between the DCB and HUS domains that we refer to as the ‘D-loop’ (Figure 3a,b, SI Appendix, Fig. S5b). In the autoinhibited cryoEM structure the D-loop interacts with helices 12 and 14 on the surface of the DCB-HUS domain, but in the predicted active state structure the D-loop interacts instead with the surfaces of helices 10 and 12. The position of the D-loop bound to the surface of the HUS domain in the autoinhibited structure, which was also observed in the isolated DCB-HUS domain crystal structure (16), appears mutually exclusive with the predicted position of the GEF domain in the active state. We therefore hypothesized that the D-loop may contribute to autoinhibition by competing with the GEF domain for binding to this surface of the Sec7 backbone.

To test this hypothesis and to characterize the relative roles of the D-loop and the HDS2- GEF domain interface in Sec7 autoinhibition we investigated the impact of corresponding mutations on *in vitro* GEF activity. We observed that the truncation of the D-loop (‘ΔD-loop’, removal of residues 444-486 in *S. cerevisiae* Sec7) resulted in a ∼8-fold increase in GEF activity towards myristoylated Arf1 on liposome membranes compared to the wild-type (Figure 4a,b, SI Appendix, Fig. S4f). In comparison, the L1376D mutation described above caused a ∼20-fold increase in GEF activity relative to the wild-type. A construct in which both of these mutations were combined exhibited a ∼50-fold increase in GEF activity relative to the wild-type (Figure 4a,b). This set of observations indicates that combining disruption of the HDS2-GEF domain interface together with truncation of the D-loop has an additive effect, and therefore both the D-loop and the HDS2-GEF domain interaction contribute to Sec7 autoinhibition.

**Figure 4.**
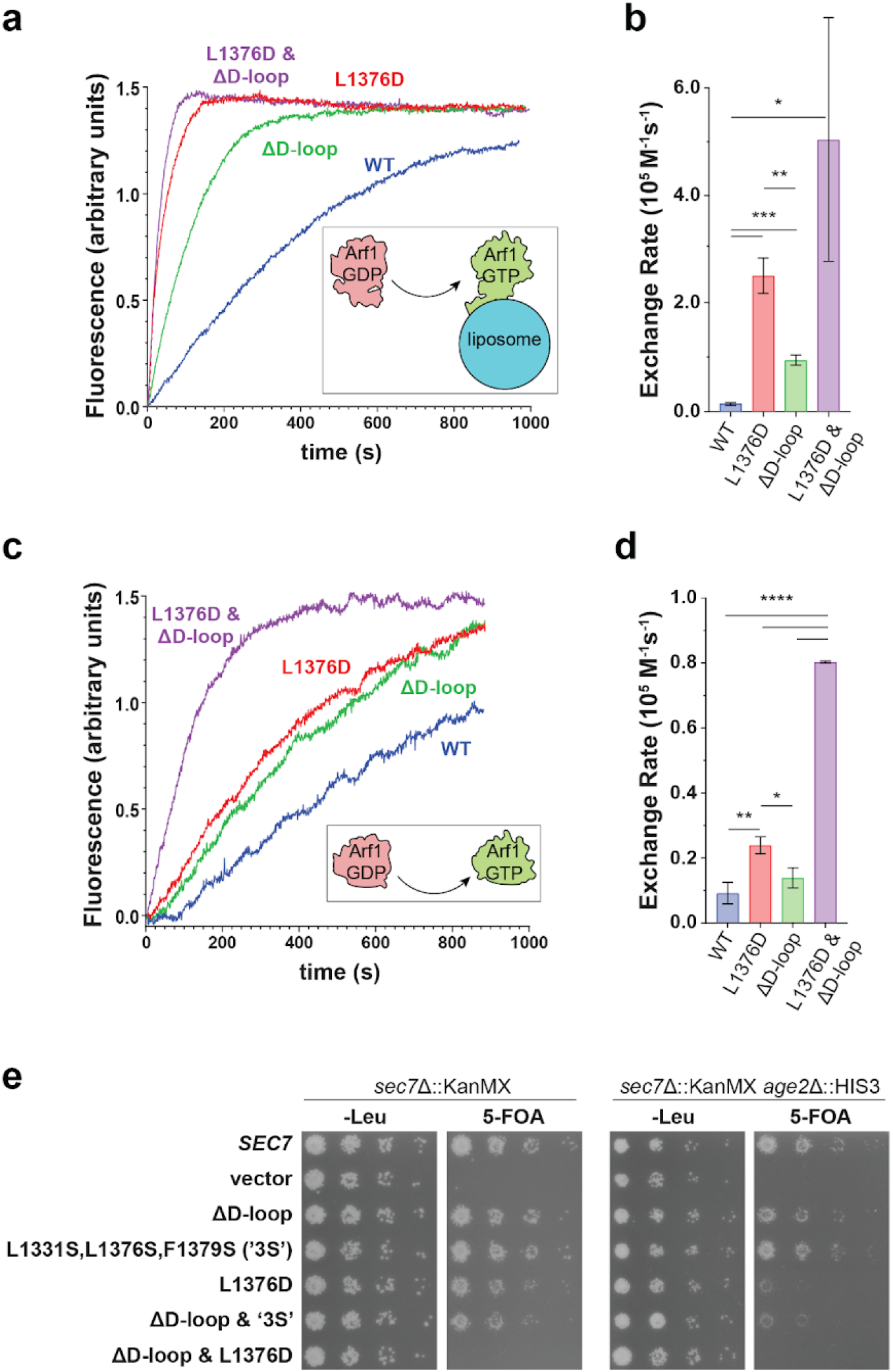
The Sec7 D-loop and GEF-HDS2 domain interface exert additive effects on autoinhibition *in vitro* and are important *in vivo*. **a-d**, GEF activity of Sec7 constructs on myristoylated Arf1 in the presence of TGN-like liposomes (a and b) or ΔN17-Arf1 in solution (c and d) monitored by native Tryptophan fluorescence. * p < 0.05, ** p < 0.01, *** p < 0.001 **e**, Complementation test (plasmid shuffling assay) testing the viability of the indicated mutants in the indicated strains.

To test whether the loss of autoinhibition caused by these mutations requires the involvement of the membrane surface, we also performed GEF assays in the absence of membranes, using the ΔN17-Arf1 substrate which does not require the presence of membranes for its activation (32, 33). Under these conditions, the three mutant Sec7 constructs displayed the same trend of hyperactivation, albeit with lower overall magnitude (Figure 4c,d). This indicates that both on membranes and in solution, the D-loop and HDS2-GEF domain interactions are important for enforcing autoinhibition of Sec7. These results, which validate the importance of the D-loop in mediating autoinhibition, also indicate the predicted structure of the Sec7 active conformation can be used to make accurate functional predictions.

We next sought to determine the relative importance of the D-loop and HDS2-GEF domain autoinhibitory interactions *in vivo*. We tested Sec7 mutants in a strain that was otherwise wild-type, as well as in a sensitized strain lacking Age2, a TGN-localized Arf-GAP (34, 35). We reasoned that decreased inactivation of Arf1 due to loss of Age2 might exacerbate the effects of excessive Arf1 activation that may arise from loss of Sec7 autoinhibition. We observed a gradient of growth phenotypes when the D-loop truncation was combined with different HDS2- GEF interface mutations (Figure 4e). We found that removal of the D-loop had no impact on growth, consistent with a previous study from our lab that first observed the D-loop bound to the surface of the isolated DCB-HUS domain crystal structure (16). The L1331S/L1376S/F1379S mutant, which perturbs multiple hydrophobic residues at the HDS2-GEF interface but does not significantly alter the charge of the HDS2 surface, also displayed no growth phenotype.

However, combining the L1331S/L1376S/F1379S and ΔD-loop mutations resulted in a mutant that exhibited a growth defect in the *age2*Δ background. The more severe L1376D mutation also resulted in a growth defect in the *age2*Δ background. Finally, combining this more severe L1376D mutation with the D-loop truncation resulted in a mutant that was inviable in both the *age2*Δ and the *AGE2* backgrounds (Figure 4e). Taken together, these results indicate that additive roles of both the D-loop and HDS2-GEF interfaces are important for Sec7 regulation *in vivo*.

Remarkably, the *in vitro* GEF activities exhibited by this series of mutants as described above (Figure 4a-d) inversely correlated with the magnitude of their *in vivo* growth defects (Figure 4e). This indicates that increasing loss of Sec7 autoinhibition *in vitro* correlates with increasingly deleterious effects on Sec7 function *in vivo*.

### Dimerization via a hydrophobic patch in the HDS4 domain

Sec7 was previously found to dimerize via the HDS4 domain (16). This domain was observed to be essential for Sec7 function *in vivo* and to have a potential role in autoinhibition (13, 14, 16). The cryoEM structure reveals the Sec7 dimerization interface is composed of 6 hydrophobic residues on each monomer (Figure 5a). With this new structural information, we sought to directly characterize the importance of dimerization by disrupting the dimer interface while keeping the remainder of the HDS4 domain intact. We therefore generated a mutant in which the six hydrophobic residues at the dimer interface are mutated to serine residues (Y1975, I1979, L1982, V1986, L1998, V2001 in S. *cerevisiae*). The purified mutant protein was monomeric *in vitro* as determined by SEC-MALS analysis (SI Appendix, Fig. S6a).

**Figure 5.**
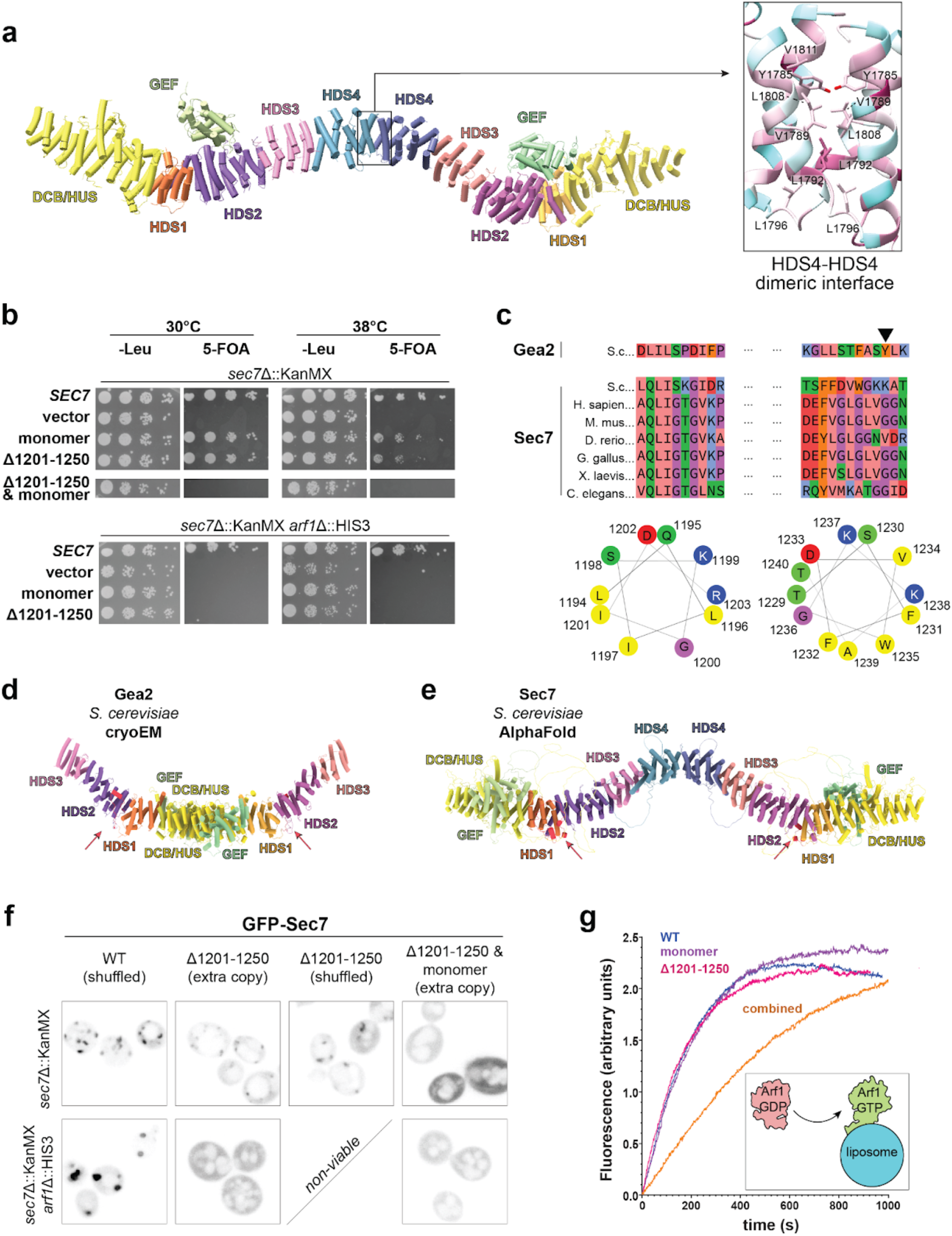
Dimerization and H-loop sequences are important for Sec7 membrane binding and localization *in vivo*. **a**, Dimeric Sec7 cryoEM structural model with boxed inset highlighting hydrophobic residues at the HDS4 dimer interface colored by conservation (magenta, highest conservation; cyan, lowest conservation). **b,** Complementation test (plasmid shuffling assay) demonstrating the impact of monomeric and H-loop (Δ1201-1250) mutations individually and in combination. **c**, Multiple sequence alignment and helical wheel projections of two short regions in the H-loop that may form amphipathic helices. Sequences shown are for *S. cerevisiae* (S. c.) Gea2 and several Sec7 homologs. In species with multiple homologs only the BIG1 homolog is included. The arrowhead indicates the Tyrosine residue that is required for Gea2 membrane binding and function (19). **d**,**e**, Gea2 cryoEM (left) and Sec7 AlphaFold (right) dimer models. The locations of the H-loops are indicated by arrows and colored red. **f**, Fluorescence microscopy of the Sec7 mutant constructs, with the indicated construct expressed as an extra copy when necessary for viability. **g**, Representative trace of Arf1 activation reactions with the indicated Sec7 construct.

To test whether dimerization is important for Sec7 function *in vivo*, we tested whether the monomeric mutant was able to complement a *sec7*Δ mutant. We observed very slight temperature sensitivity in an otherwise wild-type background but complete loss of viability in the sensitized *arf1*Δ background (Figure 5b). In contrast, removal of the entire HDS4 domain results in inviability in a wild-type *ARF1* background (16). The differences in magnitude of these growth phenotypes suggest that there may be a role for the HDS4 domain in addition to dimerization.

However we cannot rule out the possibility that loss of the HDS4 domain impacts protein folding or alternatively that the dimerization mutant does not completely disrupt dimerization *in vivo*. In either case, these results establish that dimerization is important for Sec7 function *in vivo*.

### The H-loop and dimerization are important for Sec7 membrane-binding

Recruitment of Sec7 and its metazoan homologs to the TGN has been found to involve multiple protein-protein interactions, including interactions with Rab1, Rab11, Arl1, and a positive feedback interaction with Arf1 (13, 14, 36). Several of the Sec7 regulatory domains have been implicated in TGN membrane binding (14, 16, 36, 37), but how Sec7 binds to lipids remains unclear.

The membrane binding mechanism for the related Arf-GEF Gea2 was recently identified as an amphipathic helix that is essential for membrane binding and Golgi localization (Figure 5c- e) (19). This helix is located in a loop between the HDS1 and HDS2 domains of Gea2 and is not ordered in the Gea2 cryoEM structure. We observed that Sec7 also possesses a disordered loop between the HDS1 and HDS2 domains, which we now refer to as the “H-loop”. The H-loop contains two stretches of residues which could each form amphipathic helices (Figure 5c-e).

The first lies at the C-terminal end of the HDS1 domain (residues 1194-1202 in *S. cerevisiae* Sec7), and the second is in the middle of the H-loop (residues 1229-1240 in *S. cerevisiae* Sec7) in a location similar to the membrane-binding amphipathic helix of Gea2. Sec7 homologs in other species also possess one or two amphipathic sequence regions within this loop, however we note that both differ considerably from Gea2 (Figure 5c). Furthermore, the first region corresponds to a sequence in Gea2 that is ordered and bound to the HDS2 domain far from the membrane surface (Figure 5d). In many Sec7 homologs the second region appears hydrophobic but not amphipathic. Importantly, while neither region is ordered in the Sec7 cryoEM structure, both lie near the putative membrane interacting surface in the AlphaFold predicted model (Figure 5e).

To test if the H-loop is important for the function of Sec7 *in vivo*, we generated a Sec7 construct lacking most of the H-loop to disrupt both potential amphipathic helices without disrupting the fold of the HDS1 domain. We found that this H-loop mutant (lacking residues 1201-1250) was viable in an otherwise wild-type background but was inviable in the sensitized *arf1*Δ strain (Figure 5b). However, combining the loss of the H-loop together with loss of dimerization was lethal in otherwise wild-type cells (Figure 5b).

To determine whether the H-loop is important for Golgi membrane association of Sec7 *in vivo*, we examined the localization of mutant constructs using live-cell imaging (Figure 5f). The H-loop mutant protein exhibited modestly reduced punctate intensity, either when co-expressed with wild-type Sec7 or when expressed as the only copy of Sec7 in cells. In the absence of Arf1, TGN compartments become enlarged and wild-type Sec7 is enriched on these swollen compartments. In contrast, we found that the H-loop mutant was mislocalized to the cytoplasm in *arf1Δ* cells (Figure 5f). As we were unable to image this mutant as the sole copy of Sec7 in the *arf1Δ* background due to the inviability of that genetic combination, we cannot distinguish whether the mis-localization phenotype is due to the lower overall Arf1/2 levels or competition with the endogenous copy of Sec7.

The Sec7 dimer possesses two copies of the H-loop, one in each monomer. We therefore tested the impact on localization of combining H-loop truncation with loss of dimerization. We found that a monomeric Sec7 construct also lacking the H-loop was mislocalized to the cytoplasm in otherwise wild-type cells when expressed as an extra copy (Figure 5f). This result suggests that avidity due to dimerization also plays a role in Sec7 localization to the TGN.

Taken together, these results indicate that the H-loop is important but not essential for Sec7 recruitment to Golgi compartments *in vivo*. In the absence of the H-loop it appears that the known protein-protein interactions of Sec7 and its other potential membrane-interacting regions are sufficient for Golgi localization. Yet when lacking the H-loop, dimerization is required for the essential function of Sec7 *in vivo*.

To test whether the H-loop is directly involved in lipid-membrane binding we purified H-loop (Δ1201-1250) mutant, monomeric mutant, and double-mutant constructs for *in vitro* biochemical characterization (SI Appendix, Fig. S4f). We measured the GEF activity of these purified proteins in the presence and absence of membranes. Although the monomeric and H-loop mutants behaved similarly to wild-type Sec7 under these conditions, the double-mutant construct was dramatically less active on myristoylated-Arf1 in the presence of liposomes membranes (Figure 5g, SI Appendix, Fig. S6b). In contrast, all three constructs exhibited GEF activity similar to the wild-type when activating ΔN17-Arf1 in solution (SI Appendix, Fig. S6c). To directly test the membrane-binding ability of these mutants, we performed liposome flotation experiments (25, 38). We incubated these constructs with TGN-like liposomes loaded with myristoylated Arf1-GTP, because Sec7 binds liposomes poorly in the absence of its recruiting GTPases (13, 36). We then isolated the membrane-bound proteins by flotation on a sucrose gradient (Figure 5g, SI Appendix, Fig. S7a,b). We observed that both the H-loop and monomeric mutations significantly disrupted membrane binding in this assay. These *in vitro* results correlate well with the observed Golgi localization and cell viability phenotypes, which together indicate the importance of both dimerization and the H-loop for Sec7 to activate Arf1 on the membrane surface.

### The relationship between Sec7 autoinhibition and stimulation by GTPase regulators

Sec7 GEF activity is known to be stimulated by interaction with the active forms of multiple regulatory GTPases. A positive feedback interaction with its product Arf1-GTP stabilizes Sec7 on the membrane, interactions with Arl1 and Rab1 can also recruit Sec7 to the membrane surface, and an interaction with Rab11 appears to allosterically stimulate GEF activity(13, 14, 17, 36). To characterize the relationship between autoinhibition and stimulation by these regulators we performed additional *in vitro* GEF assays with the hyperactive Sec7 constructs in the context of regulatory GTPases.

We first tested positive feedback stimulation of GEF activity by membrane-bound Arf1-GTP. We observed that not only were the basal GEF activities of the L1376D and ΔD-loop mutants significantly higher than that of the wild-type protein, their GEF activity was further stimulated by Arf1-GTP (Figure 6a, SI Appendix, Fig. S8). We then tested the impact of stimulation of GEF activity by Rab11 (yeast Ypt31) and observed that all three hyperactive Sec7 mutants were stimulated by membrane-bound Rab11-GTP to a greater extent than the wild-type (Figure 6b-d). Taken together, these results indicate that the effects on Sec7 GEF activity by regulatory GTPase stimulation and loss of autoinhibition are additive.

**Figure 6.**
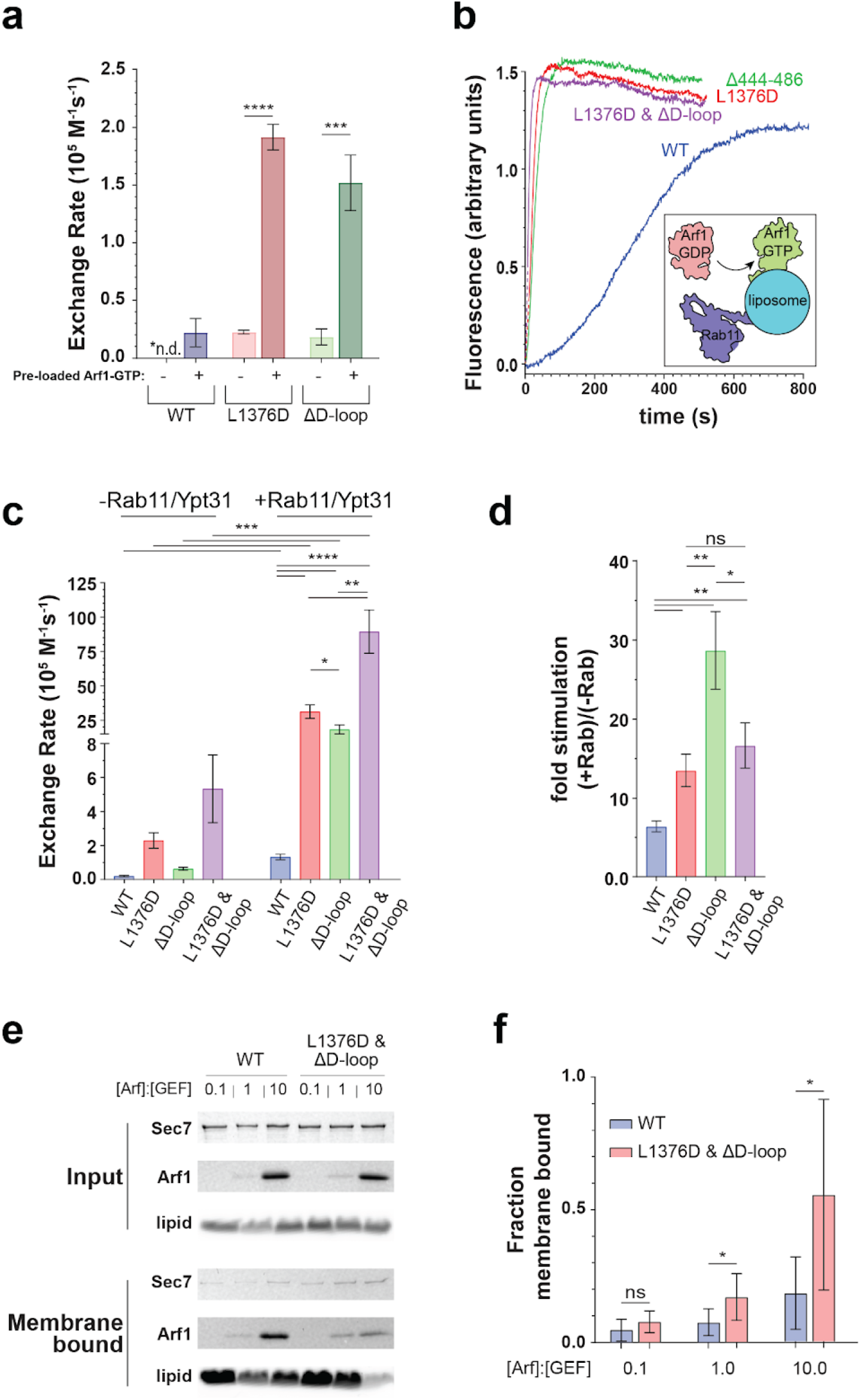
Hyperactive Sec7 is still activated by GTPases. **a**, Quantification of Arf1 GEF activity catalyzed by Sec7 constructs in the presence or absence of activated Arf1. **b**, Representative traces of Arf1 activation measured by Tryptophan fluorescence on synthetic TGN liposomes pre-loaded with Rab11/Ypt31-GTP. **c**, Quantification of triplicate measurements shown in a, (right) shown together with quantification of reactions performed without Rab11- Ypt31 (left, data also presented in Figure 4a). **d,** Ratio of exchange rates shown in b. **e**, SDS-PAGE gel of liposome floatation experiment with the indicated molar ratio of pre-loaded Arf1- GTP on liposomes. **f**, Densitometry of band intensity in e normalized by lipid recovery and input GEF. * p < 0.05, ** p < 0.01, *** p < 0.001, **** p < 0.0001

In light of these results, we wondered whether loss of Sec7 autoinhibition results in more stable membrane binding. We therefore performed a liposome flotation assay in which we titrated the amount of pre-loaded active Arf1-GTP. We found that the hyperactive L1376D/ΔD- loop double mutant was recruited to liposomes by Arf1-GTP much more robustly than was the wild-type protein (Figure 6e,f). This result indicates the active conformation of Sec7 binds membranes more stably and also suggests that membrane-bound Sec7 is more likely to adopt the active conformation.

### Model for the transition between autoinhibited and active states

To model how Sec7 binds membranes, we began with the assumption that in the fully active, membrane-bound state, the DCB-HUS domain of each monomer should be close to the membrane. This assumption is based on several pieces of evidence: The crystal structure of the human BIG1 DCB domain bound to Arl1, which is anchored to the membrane by its N-terminal amphipathic helix, indicates that the “bottom” surface of the DCB-HUS domain lies proximal to the membrane (37). Furthermore, a temperature-sensitive mutant allele (*sec7-1*), comprising a single amino acid substitution near this same surface of the DCB-HUS domain, is mislocalized to the cytoplasm at the restrictive temperature (14, 16). Finally, the similar positions of the GEF domain in both the AlphaFold Sec7 structural prediction and the cryoEM structures of Gea2 are such that membrane insertion of Arf1 during the nucleotide exchange reaction (19) would also require the same surface of the DCB-HUS domain to be located close to the membrane surface. In our autoinhibited cryoEM structure of Sec7, the overall twisted-W shape of the dimer prevents both DCB-HUS domains from being close to the membrane (SI Appendix, Fig. S9). This suggests that in addition to the occlusion of the active site caused by the interaction between the HDS2 and GEF domains, the autoinhibited conformation of Sec7 is also not compatible with stable membrane binding.

We noticed that AlphaFold predicted the Sec7 backbone to be less twisted and more curved than we observed in the cryoEM structure. To model a possible conformation of an activated dimer, we docked two copies of the AlphaFold prediction into the cryoEM density of the dimerization interface (Figure 7). In this model the overall twist of the dimer has decreased and the bend about the HDS4 domain has increased relative to the autoinhibited dimer.

**Figure 7.**
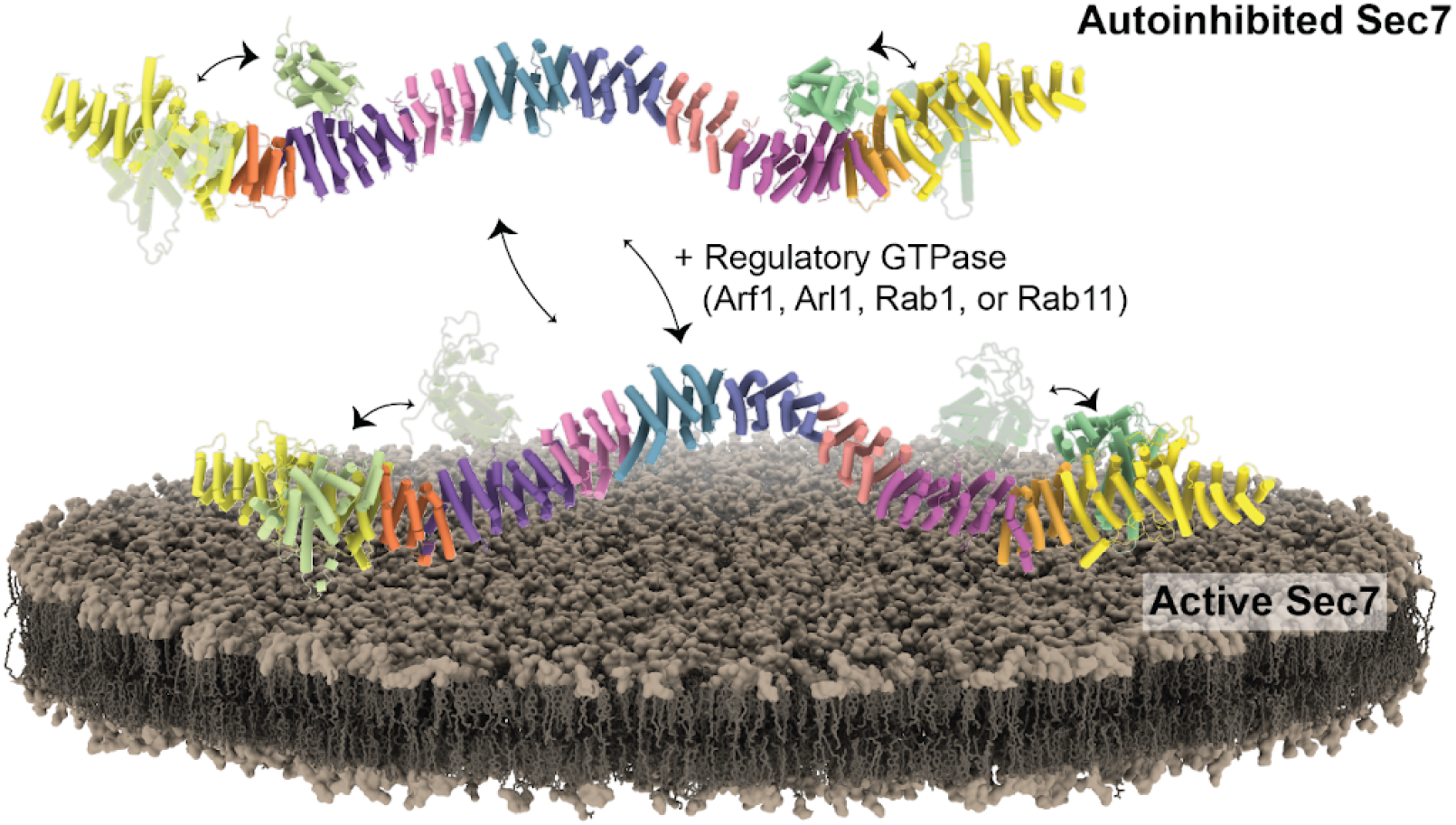
Model for Sec7 activation on the membrane. The GEF domain is in equilibrium between the two conformations. In solution, affinity for the HDS2 domain and competition with the D loop favors the GEF domain in the autoinhibited conformation. The HDS1-2 loop interacts with the membrane, which subsequently stabilizes the GEF domain in the active conformation. In the active conformation, the dimer has likely flexed to enable the DCB-HUS domain of each monomer to directly contact the membrane surface. Other factors on the membrane are then able to bind, further stabilizing the membrane bound state and/or preventing the GEF domain from switching back to the autoinhibited conformations.

Importantly, the DCB-HUS domains are closer to the membrane and both GEF domains are located such that they could insert Arf1 into the membrane during the nucleotide exchange reaction. We are therefore able to model the overall structural transitions that Sec7 needs to undergo to switch from the autoinhibited state to the active, membrane bound state (Figure 7, and SI Appendix, Video S2). In the active state each of the known interactions could be satisfied. As the H-loop extends below the backbone it may contact the membrane first and serve to loosely tether Sec7 to the surface. The conformational switch to the active state would then be promoted and stabilized by interactions with its membrane-bound GTPase regulators.

## Discussion

The experiments reported here provide detailed structural models for Sec7 in both its inhibited and active states. Our structural and functional data reveal how the arrangement and interaction of regulatory domains enables Sec7 to switch from an autoinhibited conformation in solution to an active conformation on the membrane surface. Our findings also establish important functional roles for structured loops connecting the regulatory domains.

Regulatory mechanisms governing vesicle trafficking involve a complex network of interactions. Crosstalk between GTPases, GEFs and GAPs gives spatiotemporal specificity to trafficking pathways (39–43). When coupled with positive feedback and autoinhibition, this gives regulatory pathways unique and complex properties. For example, spontaneous nucleation of Rab5 activation by the Rab5 GEF Rabex5 propagates as a wave on a supported lipid bilayer in the presence of a GAP (44). A similar dynamic behavior has also been described for the Ras GEF SOS (45). The network of interactions governing Arf1 activation by its GEF at the TGN has been documented by our group and others (13, 14, 36, 37), but the intramolecular interactions governing Sec7 regulation had not been structurally characterized prior to this work.

Autoinhibition has been documented for GEFs and GAPs of GTPases from various families including Ras, Rho, Arf, and Rab. Allosteric occlusion of the catalytic site is the typical mechanism (16, 24, 46–51), however more complex mechanisms have been suggested (52). We found that autoinhibition of Sec7 arises from an interaction between the catalytic GEF domain and the HDS2 regulatory domain. A previous study identified an allele of the *C. elegans* Sec7 homologue AGEF-1 which encodes a single residue substitution in the HDS2 domain at the GEF interface we identified (53). This allele results in enlarged late endosomes/lysosomes, and suppresses the Vul phenotype of the *let-23(sy97)* allele. This provides evidence that the autoinhibitory HDS2-GEF interaction we identified is broadly conserved and relevant to cell and developmental biological processes.

In the predicted active state, the GEF domain of Sec7 is positioned on the membrane surface by interaction with the DCB-HUS domain, similar to the cryoEM structure of the distinct Arf-GEF Gea2 (19). We found that a hyperactive mutant version of Sec7 interacted more robustly with membranes *in vitro*, suggesting that switching to the active conformation is associated with stable membrane binding. However it is also possible that Sec7 can adopt a state in which it is membrane bound yet remains autoinhibited, and this idea is supported by the fact that both Arf1-GTP and Rab11-GTP can recruit Sec7 to membranes but Rab11-GTP has a significantly stronger stimulatory effect on Sec7 GEF activity (14).

Our group previously proposed that the HDS1 and HDS4 domains have autoinhibitory roles, and that the HDS1 domain switches to an activating conformation on membranes(13, 14). In light of our new findings, several observations reconcile the previous interpretations and provide additional support for our structural model of Sec7 autoregulation. Constructs used previously to test the function of the HDS1 domain (13, 54, 55) considered the GEF-HDS1 linker to be part of the HDS1 domain. We now know that these linker residues serve to position the GEF domain relative to the DCB-HUS domain in both the Gea2 cryo-EM structure(19) and the predicted active conformation of Sec7 (SI Appendix, Fig. S5b). The importance of the GEF- HDS1 linker for Sec7 function is further supported by published mutational analysis in which a E1046A/Y1048A mutant was found to be temperature sensitive in an *arf1*Δ background (13). These residues lie at the interface between the GEF-HDS1 linker and the DCB-HUS domain.

The highly conserved ‘HUS-box’ element is also located adjacent to the GEF-HDS1 linker in these active states, explaining its important role in Arf1 activation on the membrane (17). The GEF-HDS1 linker was absent in constructs lacking the HDS1 domain, explaining why the HDS1 domain appeared to be required for robust GEF activity on membranes (13). The HDS1 domain also appeared to promote membrane binding in constructs comprising the DCB-HUS, GEF, and HDS1 domains (13, 17, 56). However, these HDS1-containing constructs used previously were truncated at residue 1220 and therefore include the conserved amphipathic helix present at the beginning of the H-loop, which we have found is important for membrane binding. Taken together these findings also explain why inclusion of the HDS1 domain stimulated GEF activity of Sec7 constructs on membranes but not in solution (13).

It is unclear why removal of the HDS4 domain resulted in loss of autoinhibition to a similar extent as loss of the HDS1-4 domains (14), but we surmise that truncation of the entire HDS4 domain may have perturbed the ability of the HDS2 domain to interact with the GEF domain. Our cryoEM structure of Sec7 enabled us to disrupt dimerization more surgically, resulting in our new finding that an important function of dimerization is to provide avidity for membrane binding. Disruption of dimerization also enabled us to uncover the important role of the H-loop in Sec7 membrane binding.

It remains unresolved why Sec7 evolved to be regulated by autoinhibition yet the related Arf-GEFs Gea1/2 did not. It is therefore interesting to consider the physiological consequences of Sec7 hyperactivation *in vivo*. Our results indicate that Sec7 hyperactivation above some threshold is deleterious, but the precise impact this has on trafficking and why partial hyperactivation is tolerated remains to be determined. In contrast to a canonical signaling output, in which hyperactivation of a GEF is expected to drive hyperactivation of its GTPase signaling pathway, the loss of Sec7 autoinhibition may not necessarily result in increased Arf1 activation at the TGN. For example, loss of Sec7 autoinhibition could potentially cause ectopic activation of Arf1 at other cellular locations, resulting in reduced Arf1 activation at the TGN. The *C. elegans* HDS2 domain mutant which we predict to disrupt autoinhibition results in a loss-of-function, rather than gain-of-function, phenotype for the Sec7 homolog AGEF-1 (53). Similarly, we found that reduction of the Arf1/2 level in cells exacerbated the hyperactive mutant growth phenotypes, rather than rescued as would be expected in a simple signaling cascade. Further studies, using the structural information generated in this study, are needed to characterize how Sec7 hyperactivation impacts Golgi trafficking.

Although we validated a predicted model for the active conformation of Sec7, the experimental structure of Sec7 in its active state on a membrane will be important to determine. There may be multiple paths between inactive/freely-diffusing and active/membrane-bound Sec7, and more than one active conformation. Future studies are needed to determine precisely how Sec7 interacts with its regulatory GTPases on the membrane surface.

## Methods

### Cloning and strain modification

All constructs (SI Appendix, Table S1) were created using NEB HiFi gibson master mix, and are full-length, N-terminally tagged, and otherwise only modified as indicated. All constructs were verified by sequencing. Yeast strains (SI Appendix, Table S2) and plasmid transformants were generated by standard LiOAc (*S. cerevisiae*) or electroporation (*Pichia pastoris*) of appropriate DNA constructs (see SI Appendix).

### Protein purification

See SI Appendix for full purification details, briefly *Komagataella pastoris* carrying Sec7 constructs under the AOX1 promoter were cultured with autoinduction media as described (57), collected by centrifugation after 48hrs, and lysed in a Spex cryogenic mill (6875D). After addition of lysis buffer and clarification, lysate was loaded onto NiNTA resin, washed and eluted.

*T. terrestris* Sec7 for cryoEM was purified using a HisTrap column (Cytiva Cat. No. 29051021), using an AKTA Pure for wash and elution. Batch NiNTA resin (Thermo Cat. No. 88221) that had been fragmented by sonication was found to have the highest capacity for *S. cerevisiae* Sec7 constructs, after loading and washing by centrifugation. Elution of *S. cerevisiae* Sec7 was performed by protease cleavage, and further purified by size exclusion chromatography on a Superose 6 Increase 10/300 GL column (Cytiva Cat. No. 29091596). Elution fractions were checked by SDS-PAGE, pooled, concentrated, flash frozen in liquid nitrogen, and stored at - 80°C for later use.

ΔN17-Arf1, myristoylated Arf1, and prenylated-Ypt31/GDI complex were purified as previously described (13, 25, 58).

### Preparation of fragmented affinity resin

In order to increase the surface area of the Ni-NTA resin accessible to the large Sec7 molecule, the resin was resuspended in water to make a 20% slurry, then sonicated at 80% power for three minutes with a macro-tip, (cycles: 20s on / 10s off). The extent of fragmentation was checked under a light microscope (4x magnification) to verify the integrity of most agarose beads had been disrupted. Following sonication we equilibrated the fragmented resin 5x with buffer at a lower speed (1000 rpm) to remove resin fines.

### CryoEM sample preparation, data collection, and processing

Grids (Quantifoil R1.2/1.3) were glow discharged in a Pelco easyGlow for 60s, 10 mA, under a 8:2 argon:oxygen gas mix. Fluorinated fos-choline 8 (Anatrace, cat# F300F) was added to *T. terrestris* Sec7 to a final concentration of 2 mM. 3 µl of this protein solution was applied to the grid, which was then blotted for 3 s and immediately plunged into liquid ethane using a Vitrobot Mark IV. CryoEM data was collected at 63 kX nominal magnification on a Talos Arctica operating at 200 kV equipped with a K3 detector and a BioQuantum energy filter. In total, 4,474 movies were collected, each with 129 frames and a total dose of 55.6 e/A2. Automated data collection was done with SerialEM (59) using aberration-free image shift to collect multiple exposures per stage shift.

Standard cryoEM data processing tools (MotionCor2, GCTF, CryoSPARC, Relion 3.1) (60–63) were used to correct beam induced motion, estimate CTF parameters, pick, sort, and symmetry expand particles, and refine and reconstruct the final maps. The published crystal structure of the DCB-HUS domain and AlphaFold prediction (16, 18) were used for guidance with *de novo* building in the few regions with poor side chain density. Atomic models and composite maps were generated, refined, and validated using Real Space Refine (64) and Phenix Combine Maps in Phenix (65–67). For cryoDRGN analysis, we used TOPAZ to increase the likelihood of rare particles (68) (this did not improve resolution of the monomer) and aligned sorted particles on a monomer without subtraction for input into cryoDRGN (20). See Tables 1 and 2, and SI Appendix, Figs. S1, S2, and S3.

### Yeast complementation tests (plasmid shuffling assay)

Yeast expression plasmids with Sec7 constructs were transformed into the indicated strain: CFY409 (*sec7*Δ), CFY863 (*sec7*Δ*arf1*Δ), or CFY4969 (*sec7*Δ*age2*Δ) as described above and in SI Appendix. Single colonies were collected and normalized for cell density, then serial diluted and pinned on indicated media. Plates were incubated for three days at the indicated temperature (30 °C if not otherwise specified) before imaging.

### Fluorescence microscopy

Cells carrying the indicated constructs were grown to an OD of 0.6 before imaging. Cells were allowed to settle on a coverslip dish (MatTek) for 10 min, and washed with fresh media just prior to imaging. See SI Appendix for imaging details. Exposure and laser power were adjusted according to intensity, and were kept the same for all specimens being compared in an experiment. The brightness/intensity was equivalently adjusted across all images in an experiment using ImageJ.

### Liposome preparation

Synthetic TGN liposomes were prepared (25) and used for liposome floatation (38). See SI Appendix, Table S3 for their composition, and SI Appendix for a detailed protocol. Briefly, after extrusion 250 µM liposomes were loaded with Arf1-GMPPNP by EDTA exchange. Sec7 constructs were then added to a final concentration of 550 nM, incubated at room temperature for 1 hour, and separated from unbound protein by ultracentrifugation under a sucrose gradient. Protein and lipid recovery was analyzed by SDS-PAGE (12% acrylamide gel).

### In vitro Arf activation (GEF) assay

GEF activity was determined by measuring the native Tryptophan fluorescence of Arf1, as described previously (25) and detailed in SI Appendix. Briefly, to a final volume of 150 µl, 200 µM 100 nm TGN liposomes, Sec7 construct (with concentration as detailed below), 1 µM myristoylated Arf1, and 200 µM GTP were added to a quartz cuvette. For Figures 2 and 4, GEF was added to a final concentration of 100 nM. For Figure 6b, 20 nM GEF was added. For Figure 6a, 60 nM GEF was added. For Figure 5g and SI Appendix, Fig. S6c, 200 nM GEF was added. The order listed is the order components were added, except for reactions with preloaded Ypt31. For these reactions, 500 nM prenylated Ypt31 was loaded onto liposomes by EDTA exchange of GMPPNP. The presence of GMPPNP in the liposome mixture dictated that Arf1 was added last. Tryptophan fluorescence (297.5 nm excitation, 340 nm emission) was measured using a fluorometer, and curves were fitted in GraphPad PRISM 10 using a nonlinear regression (one phase association).

### Statistical analysis

All assays were performed in triplicate, and statistical analysis was performed using GraphPad PRISM 10. GEF assay comparisons were analyzed by unpaired parametric t-test. Liposome floatation comparisons were analyzed by paired ratio t-test.

### Data Deposition

CryoEM maps have been deposited in the EMDB (EMD-42135, EMD-42182, and EMD- 42183) and structure coordinates have been deposited in the RCSB PDB (8UCQ).

## ACKNOWLEDGEMENTS

We acknowledge the Cornell Center for Materials Research (CCMR), especially K. Spoth and M. Silvestry-Ramos, for access and support of electron microscopy sample preparation and data collection. The CCMR is supported by NSF grant DMR-1719875. We thank members of the Fromme lab for helpful advice and discussions, especially J. Ryan Feathers and Saket Bagde, and Maria Font from Holger Sondermann’s laboratory for assistance with MALS data collection and analysis. This study was funded by NIH grant R35GM136258 to J.C.F.

## AUTHOR CONTRIBUTIONS

B.A.B performed all reported experiments and data analysis. B.C.R. and S.L.H. performed extensive protein crystallization trials and preliminary truncation analyses. J.C.F. supervised the project and obtained funding. B.A.B. and J.C.F. wrote the manuscript.

## Declaration of Interests

The authors declare no competing interests.

## SI Appendix

**SI Appendix, Figure S1.**
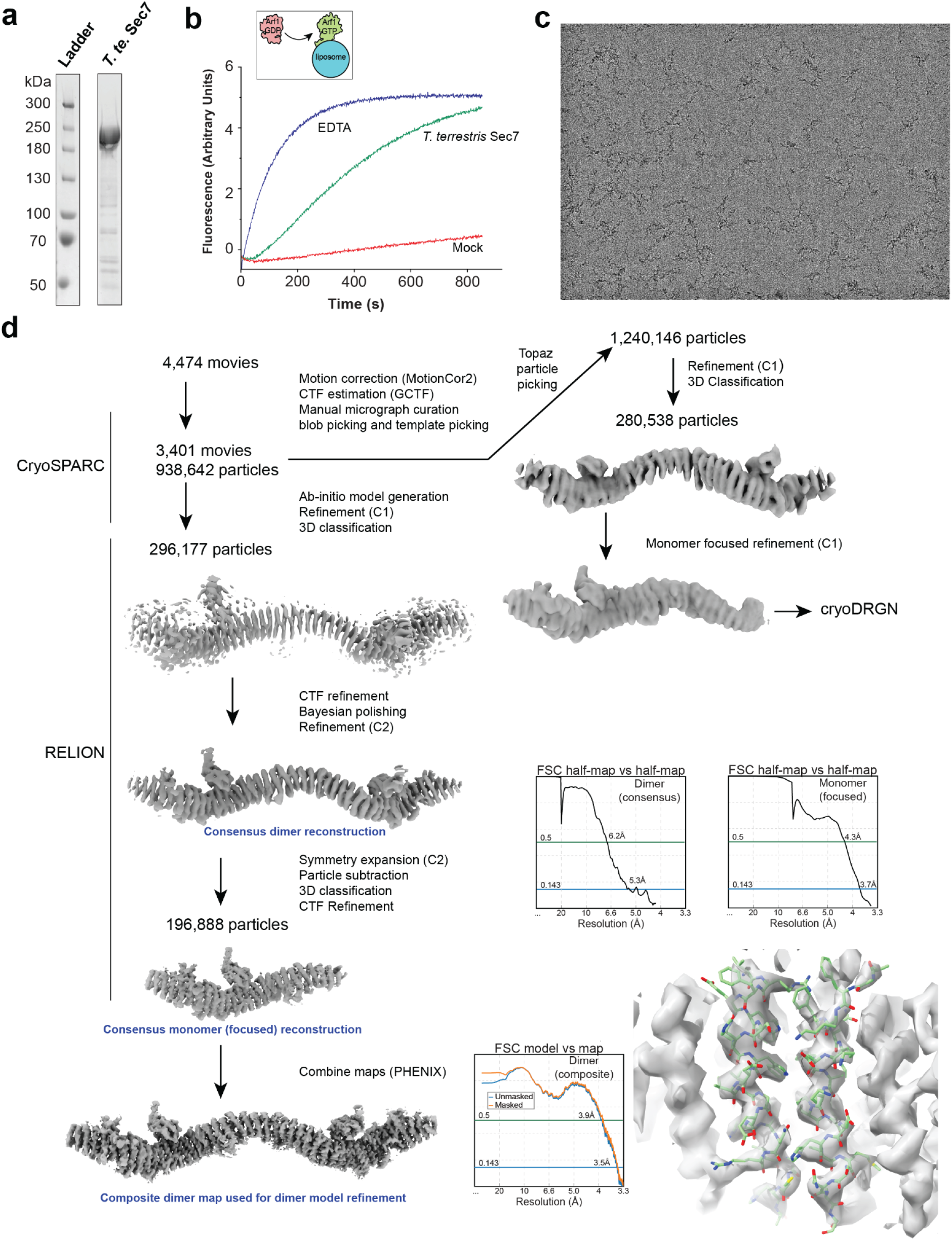
*T. terrestris* Sec7 purification and CryoEM data processing workflow. **a**, SDS-PAGE of purified *T. terrestris* Sec7 expressed in *P. pastoris*. **b**, Tryptophan fluorescence GEF activity assay demonstrating that the purified *T. terrestris* Sec7 construct is capable of myristoylated-Arf1 activation in the presence of TGN liposomes (see Methods). Note that the observed rate and sigmoidal shape of the curve are consistent with autoinhibitory behavior previously determined for *S. cerevisiae* Sec7 (1). **c**, Representative cryoEM micrograph. **d**, CryoEM data processing workflow.

**SI Appendix, Figure S2.**
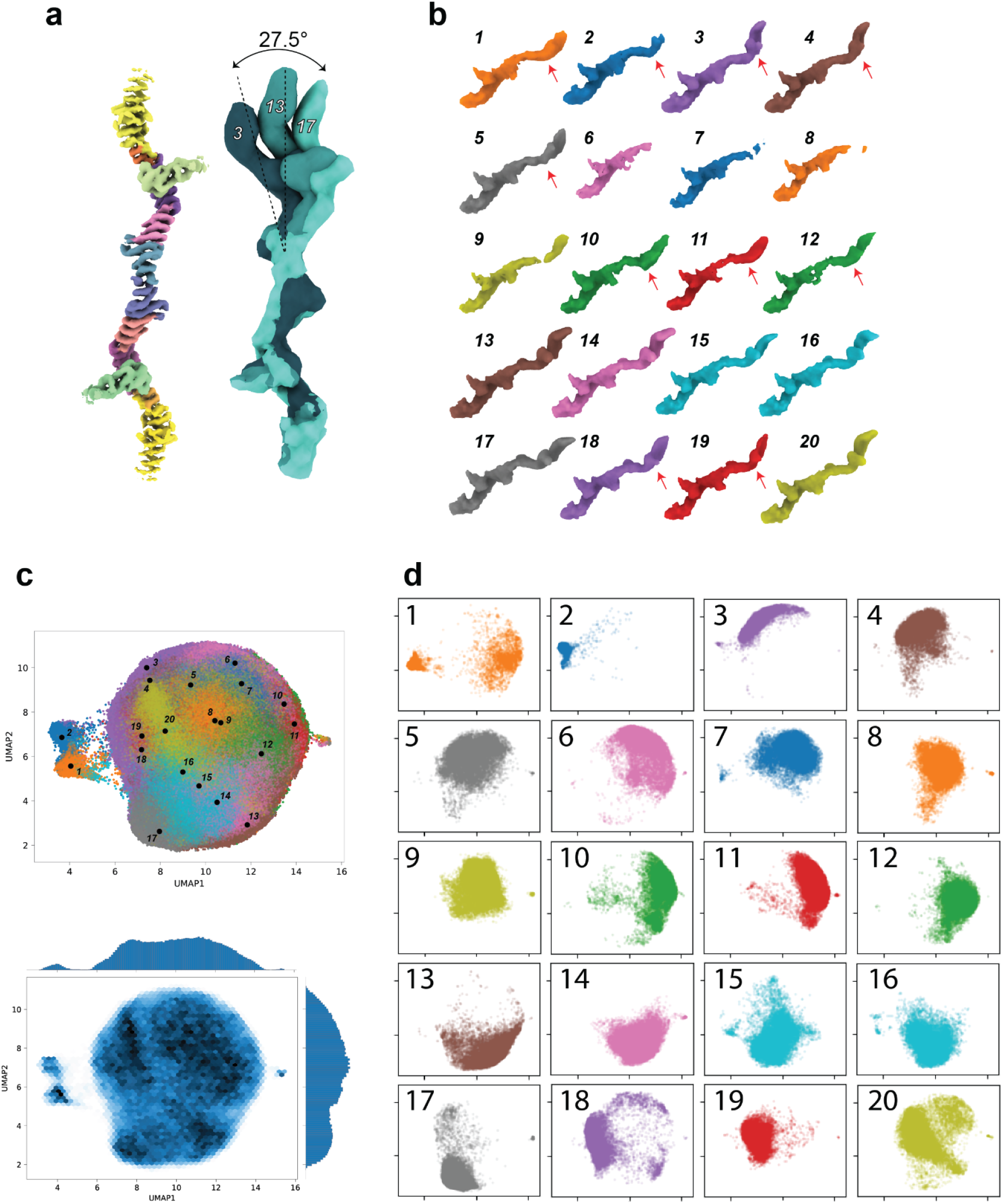
Heterogeneous Reconstruction and UMAP analysis by cryoDRGN. Dimer particles were focus-refined on a monomer (without symmetry expansion or subtraction) and used to train a cryoDRGN variability model (2). **a**, Cryo EM map of the Sec7 dimer reconstruction (left), and three example cryoDRGN generated maps highlighting the flexibility of the Sec7 dimer. **b**, All maps generated by cryoDRGN. Maps 6-10 significantly lack density for a second monomer, perhaps due to flexibility. Red arrows indicate maps with little/no density for the GEF domain bound to HDS2. We note that this only occurs in the monomer that was not focused on during refinement, and therefore could be artifactual. **c**, Plots of UMAP analysis colored by reconstruction weight of the maps in b (top) or by density of particles (bottom). **d**, Isolated UMAP distribution plots of each map in b.

**SI Appendix, Figure S3.**
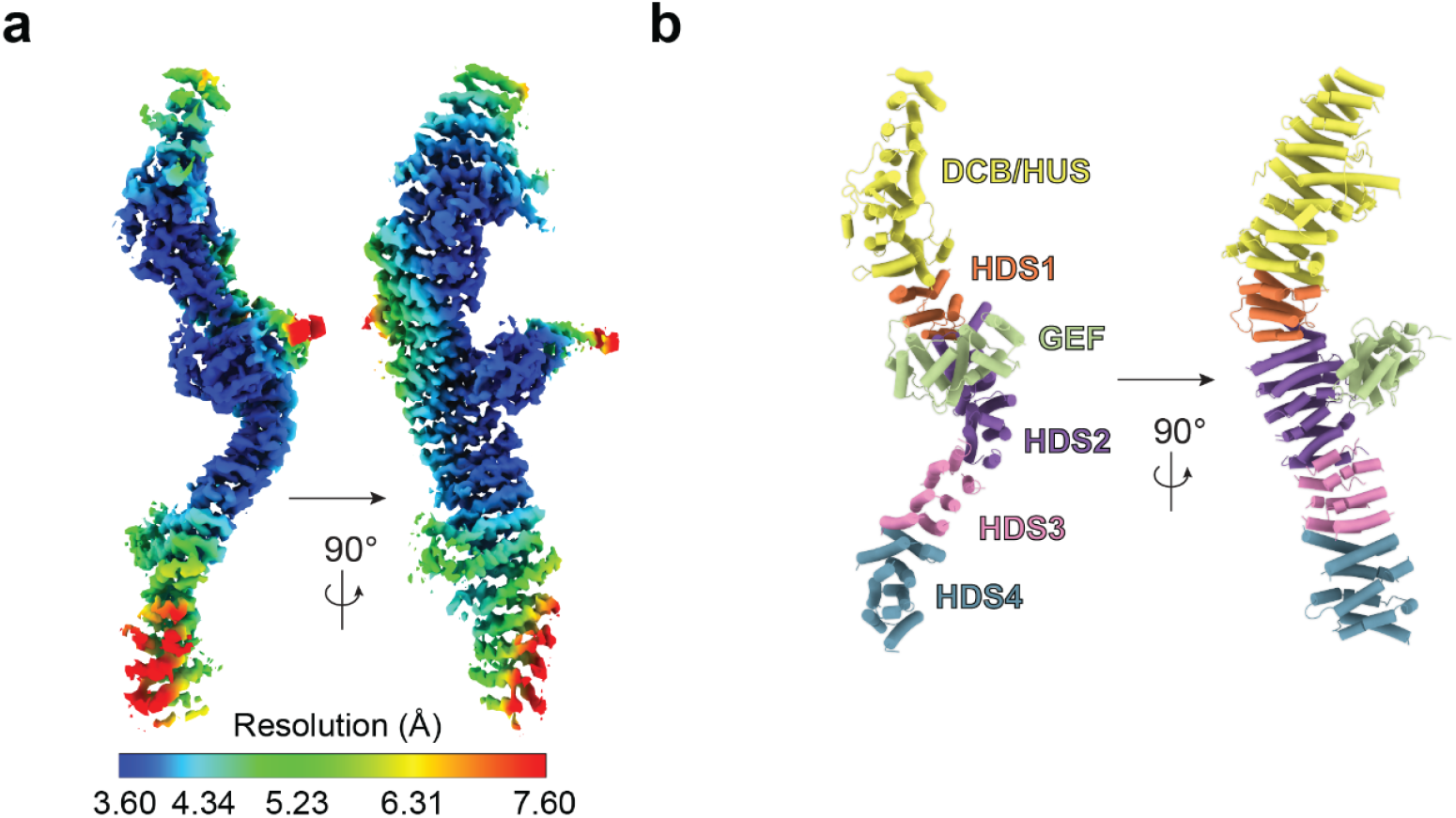
Focused refinement of a Sec7 monomer. **a**, Map generated by focused refinement of an entire single monomer using symmetry expanded particles gave the highest resolution, 3.7 Å overall. Local resolution determined using RELION (3) was used to color the map as indicated. **b**, Model built and refined using the map in a, colored by domains as indicated.

**SI Appendix, Figure S4.**
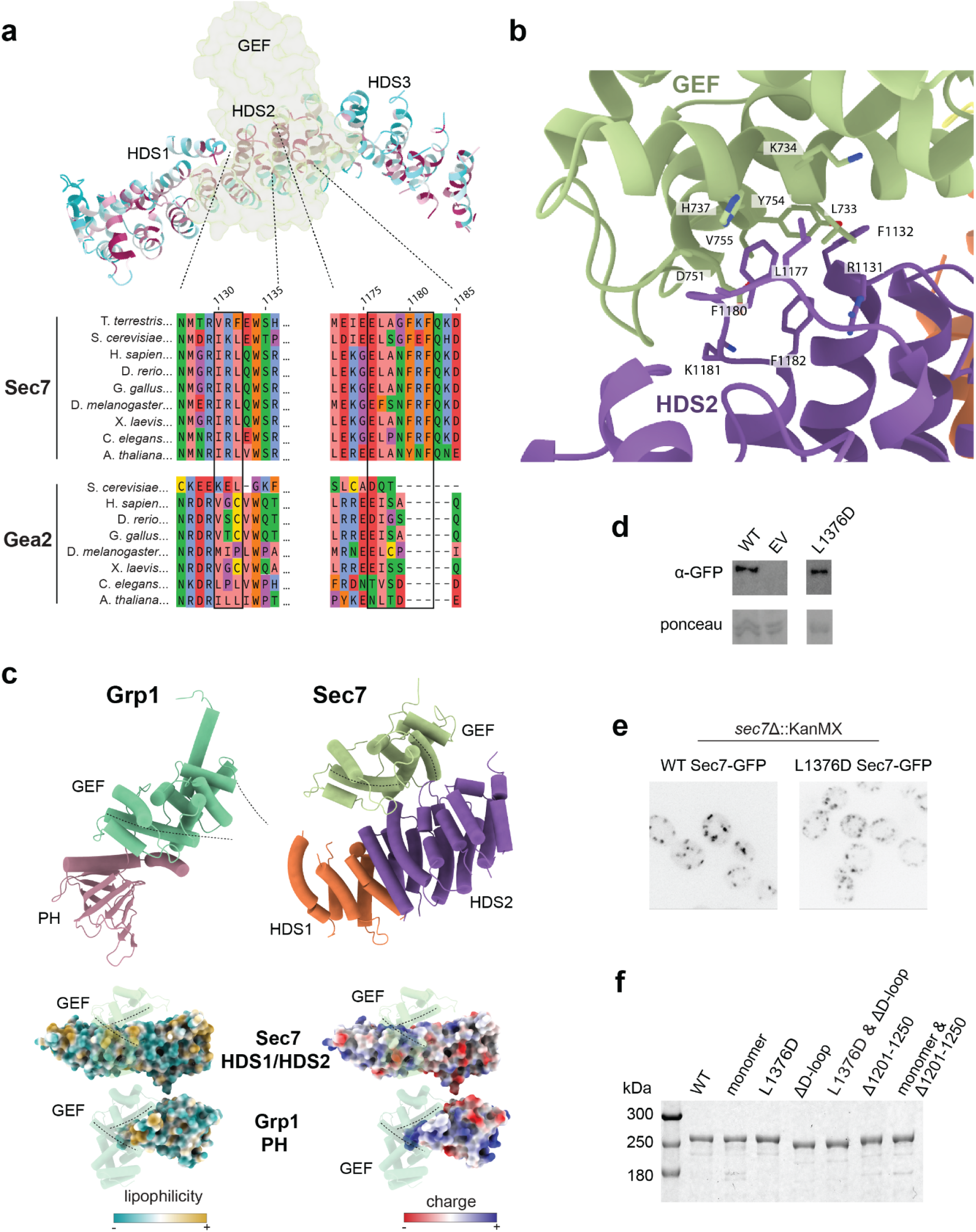
Detailed view of HDS2 interface. **a**, Multiple sequence alignment of the HDS2 interface in Sec7 and Gea2 homologs. Boxed regions indicate the surface exposed loops that contact the GEF domain. **b**, Key residues of the HDS2 interface are shown. **c**, Comparison with the autoinhibited Grp1 crystal structure. Grp1 autoinhibition involves a basic patch in the PH domain that interacts with its GEF domain in a distinct manner from that of the Sec7 HDS2-GEF domain interaction. **d**, Western blot for α-GFP-Sec7, showing the L1376D construct is expressed similarly to WT after shuffling. **e**, Microscopy of GFP-Sec7 constructs after shuffling in *sec7*Δ::KanMX strain background. **f**, Example SDS-PAGE gel of purified protein constructs used in assays shown in Fig. 2d,e and Fig. 4a-d.

**SI Appendix, Figure S5.**
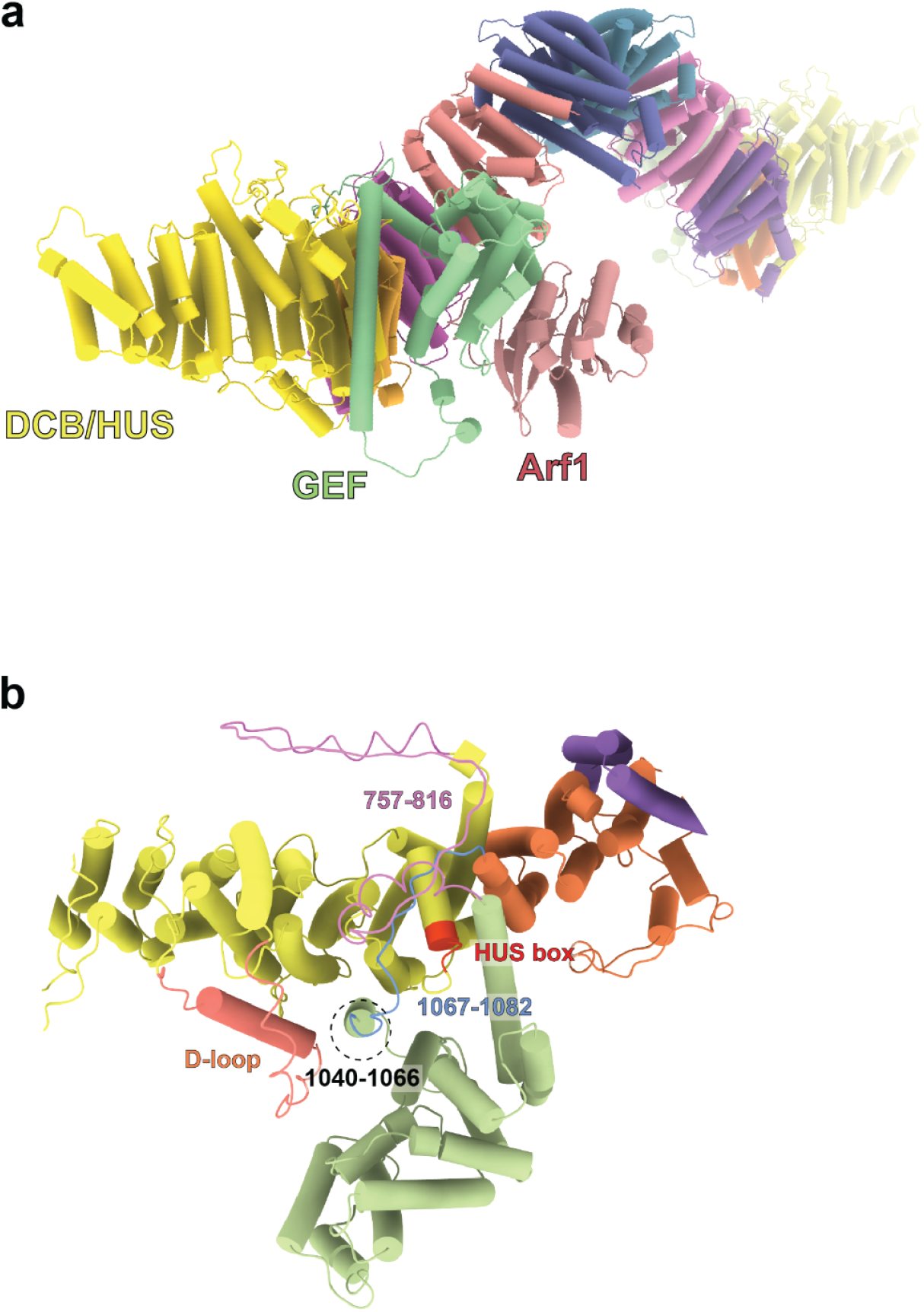
Predicted structural model of Sec7 in an active conformation. **a,** AlphaFold prediction with Arf1 bound to the GEF domain as in the Gea2-Arf1 cryoEM structure (4) shows the catalytic surface of the GEF domain is accessible in this conformation. **b**, Positions of the GEF linkers (757-816 and 1067-1082 in S. cerevisiae) surround the HUS box in the AlphaFold prediction, and physically connect the GEF domain to the activating surface of the DCB/HUS domains. Note residues 1040-1082 correspond to the GEF-HDS1 linker identified in the Gea2 cryoEM structure (4).

**SI Appendix, Figure S6.**
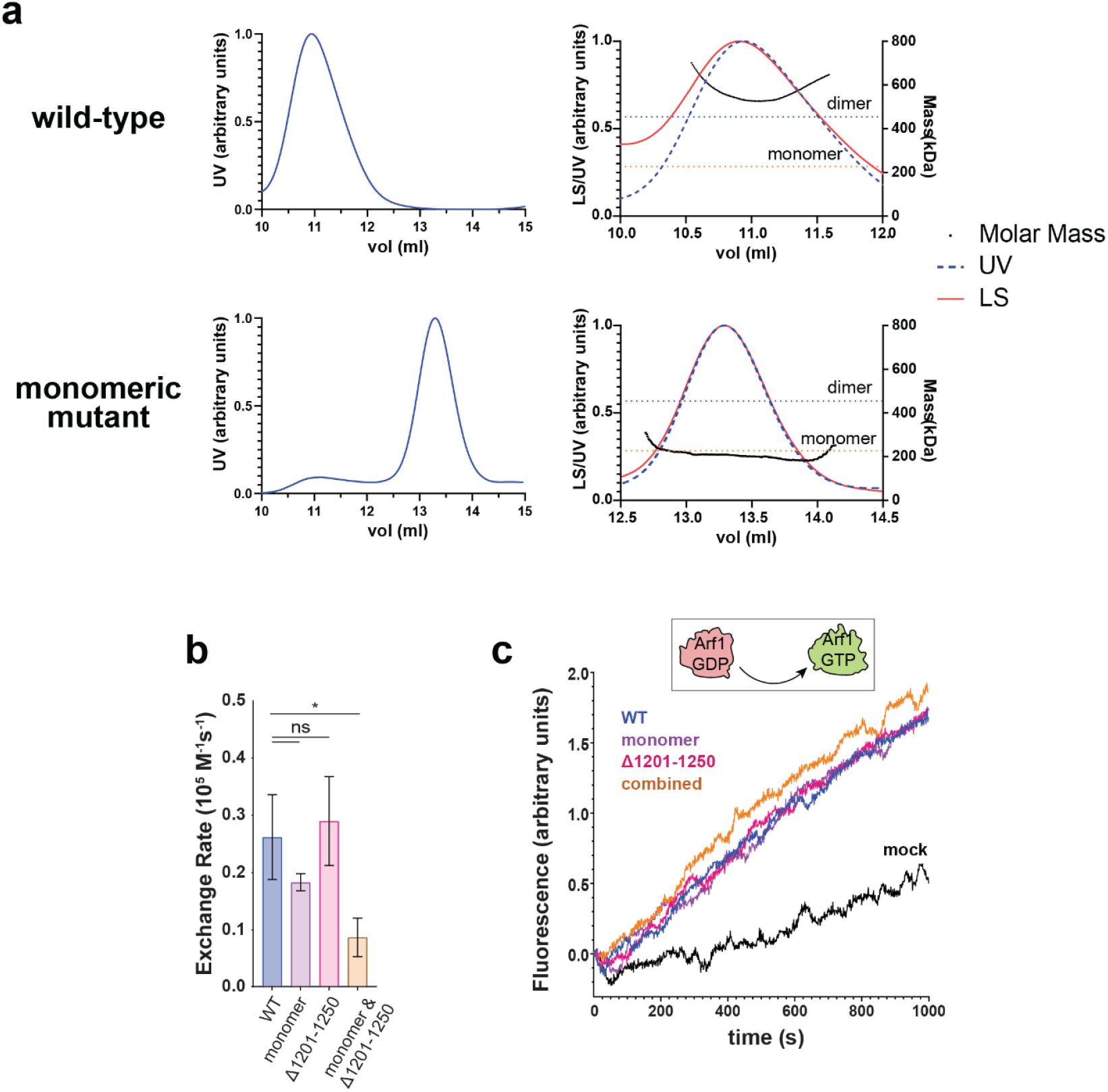
SEC/MALS of monomeric Sec7 construct and membrane binding. **a**, Purified monomeric Sec7 eluted ∼2.5 ml later than WT Sec7 over a Superose 6 10/300 column (left), and was verified by MALS to be monomeric (right). **b**, Quantification of reaction rate constants for triplicate measurements as reported in Fig. 5g. n.s.: not significant, * p < 0.05 **c**, Representative traces of ΔN17-Arf1 exchange catalyzed by the indicated GEF constructs.

**SI Appendix, Figure S7.**
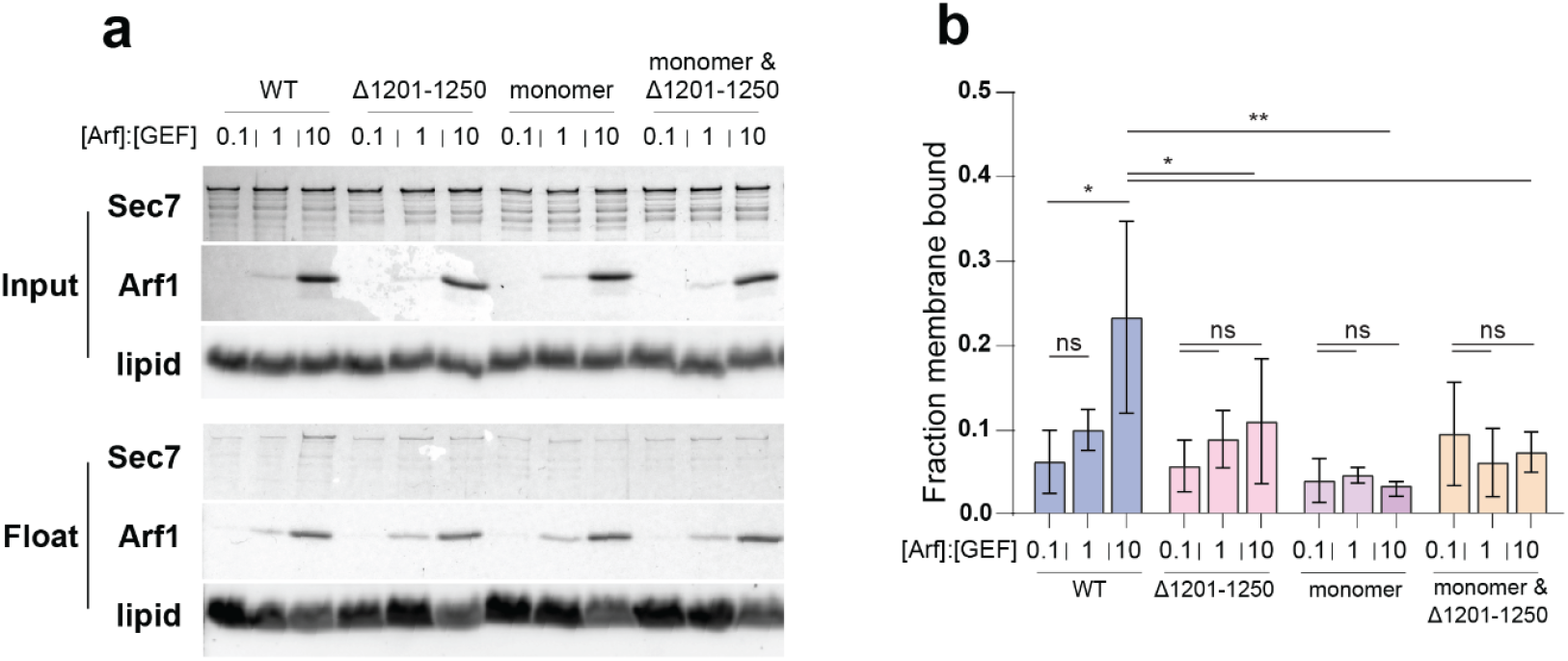
Dimerization and membrane binding. **a**, Representative SDS-PAGE of membrane float assay. **b**, Quantification of triplicate measurements in a. n.s.: not significant, * p < 0.05, * p < 0.01

**SI Appendix, Figure S8.**
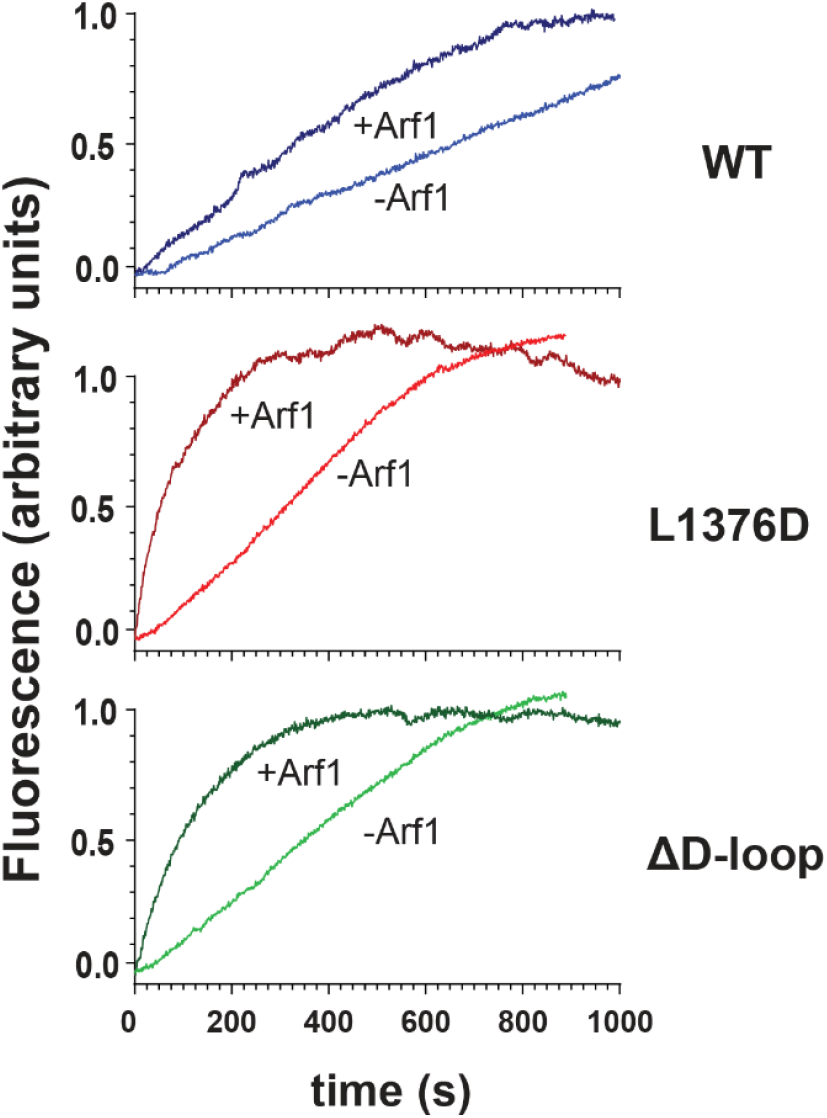
Raw data for Fig. 6d. Representative trace of GEF assay shown in Fig 6d.

**SI Appendix, Figure S9.**
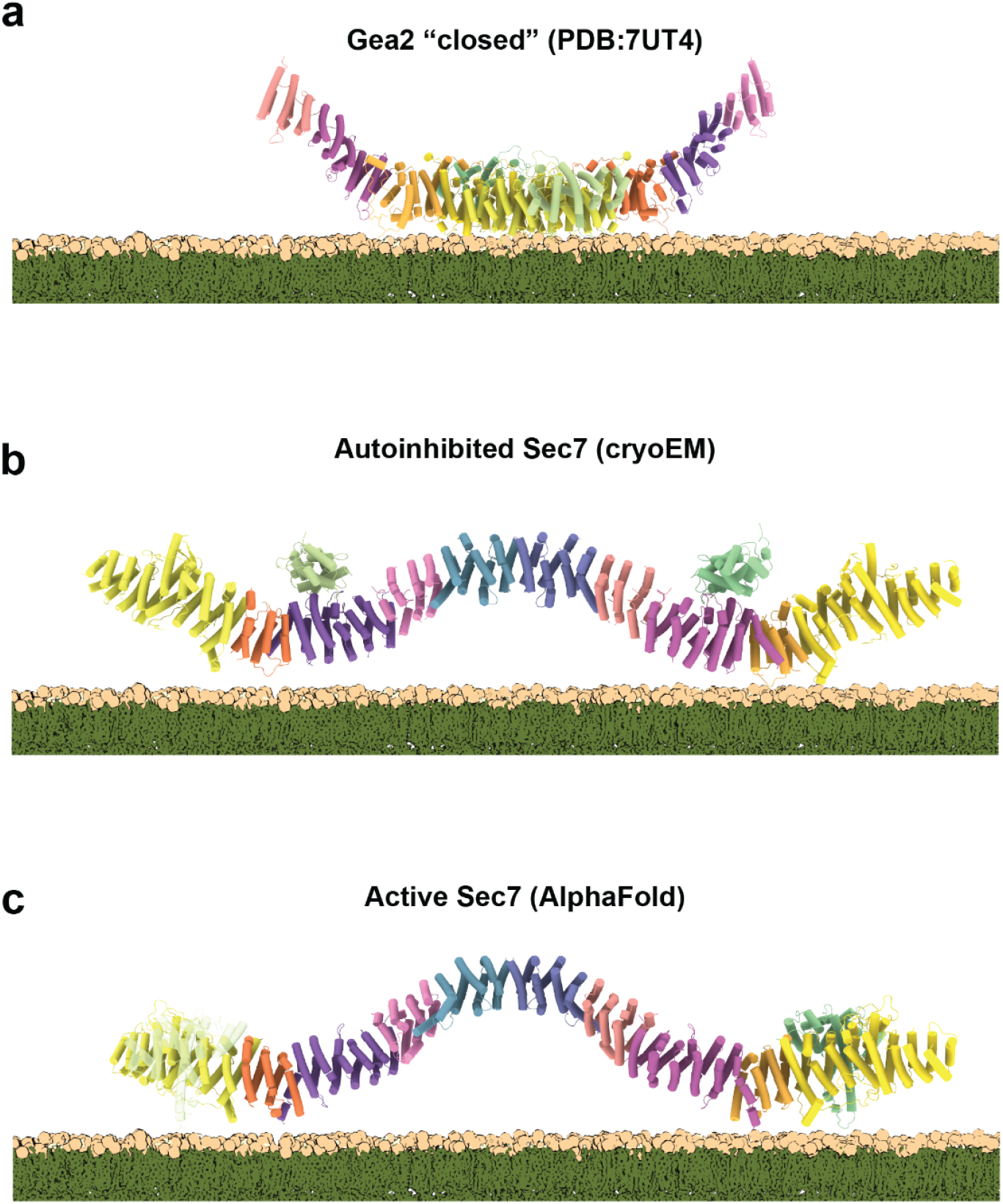
The autoinhibited conformation of Sec7 is not compatible with stable membrane binding. Comparison of Gea2 cryoEM (**a**), autoinhibited Sec7 cryoEM (**b**), and AlphaFold-predicted Sec7 (**c**) structures on the membrane surface.

**SI Appendix, Table S1.**
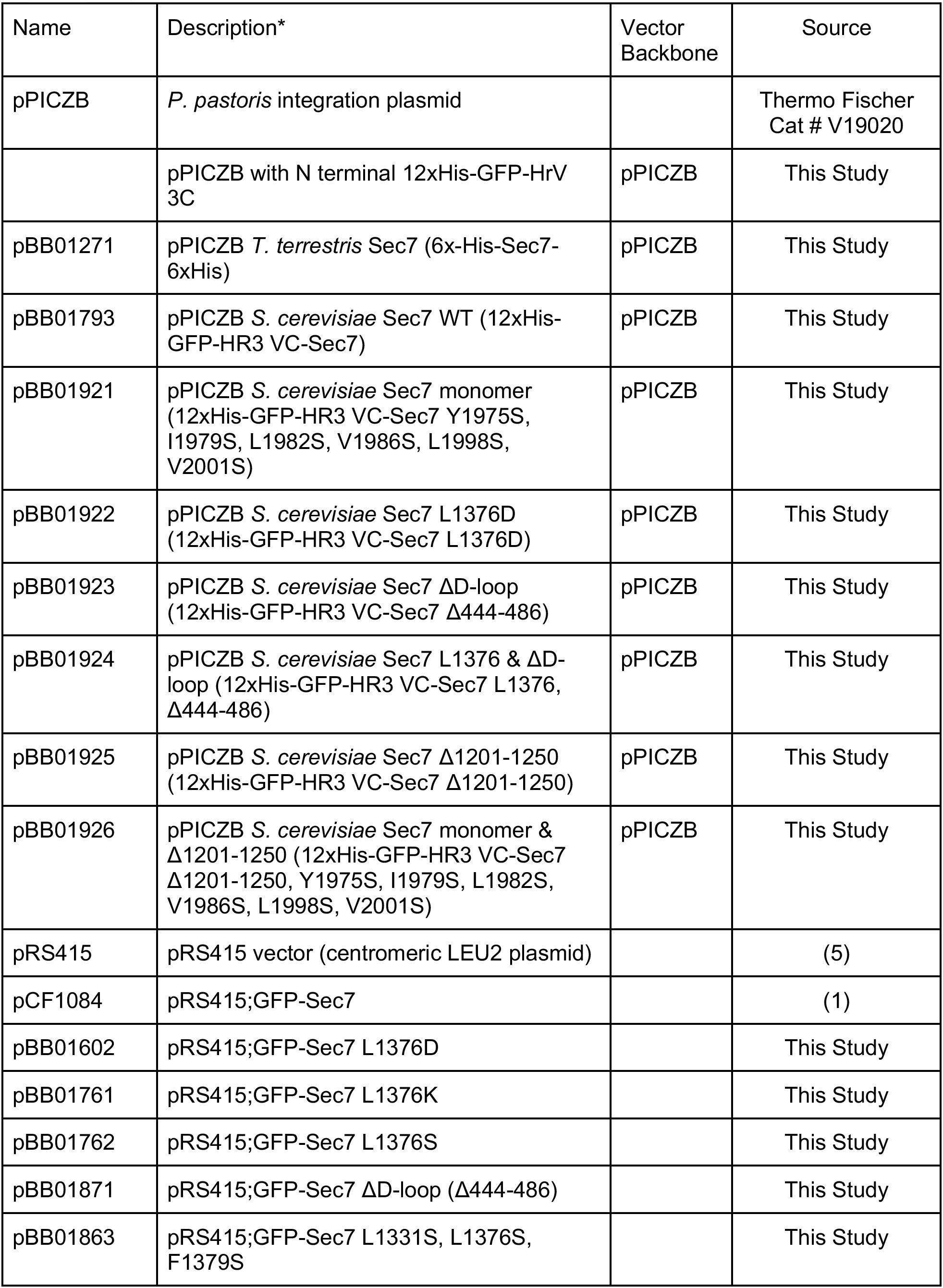
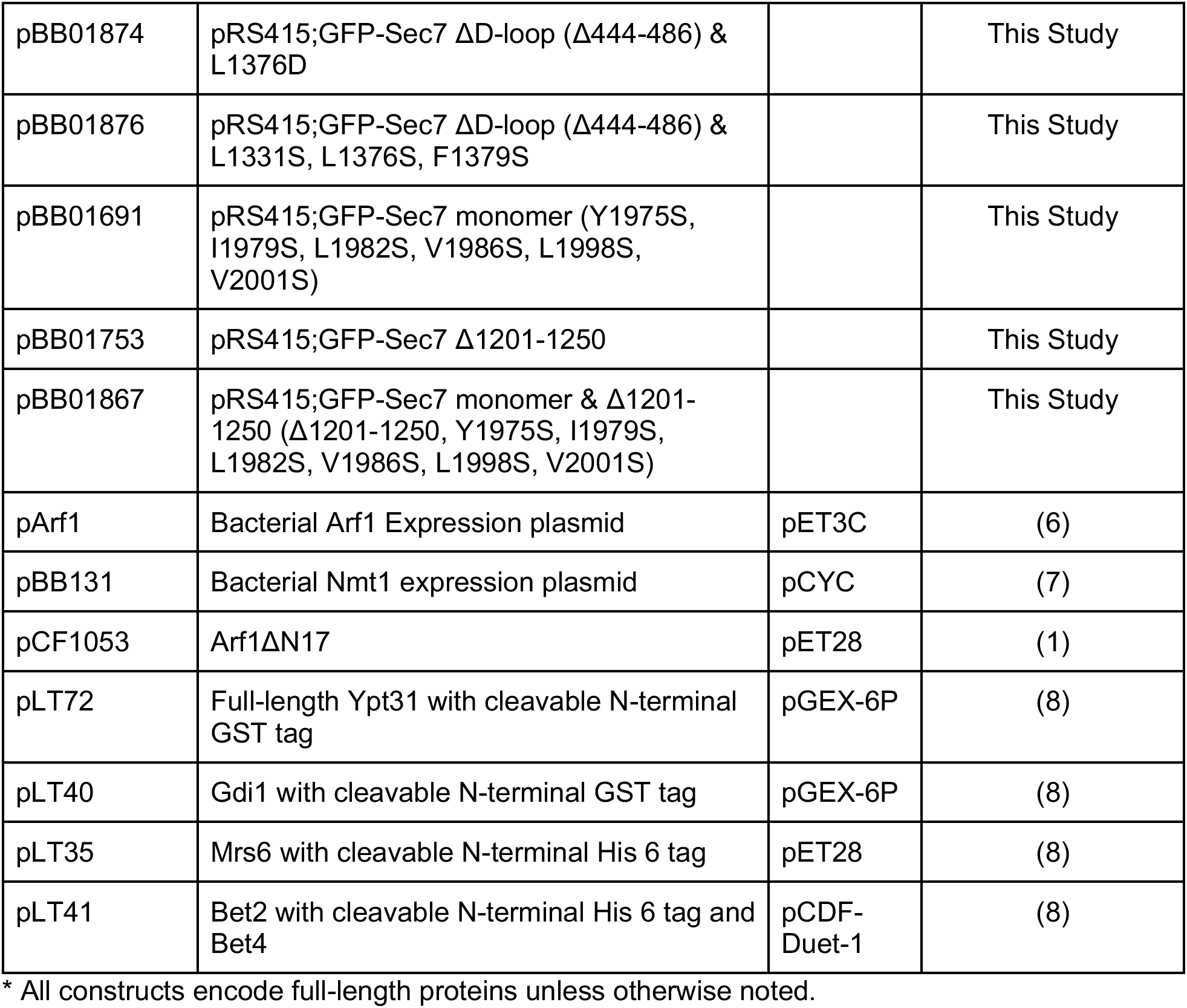
Plasmids.

**SI Appendix, Table S2.**
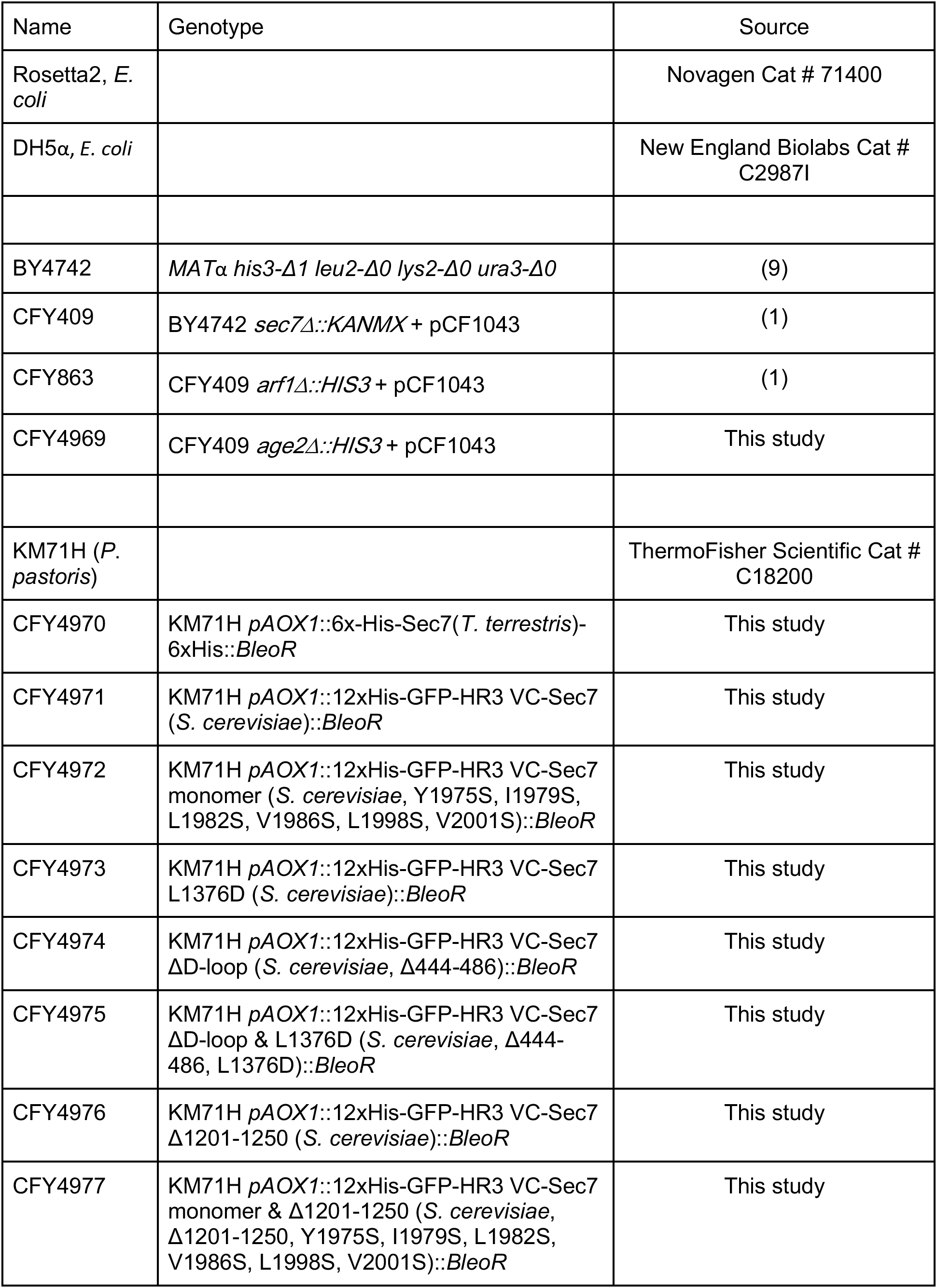
Bacterial and Yeast Strains.

**SI Appendix, Table S3.** TGN lipid mix.

| <u>Lipid</u> | <u>mol%</u> |
| --- | --- |
| DOPC | 24 |
| POPC | 6 |
| DOPE | 7 |
| POPE | 3 |
| DOPS | 1 |
| POPS | 2 |
| DOPA | 1 |
| POPA | 2 |
| Liver or Soy PI | 30 |
| PI(4)P | 1 |
| CDP-DAG | 2 |
| PO-DAG | 4 |
| DO-DAG | 2 |
| Ceramide (C18) | 5 |
| Cholesterol | 10 |
| DiR | 1 |

## SI Appendix, Video Legends

### SI Appendix, Video S1

Video of the Sec7 dimer cryoEM structure. The presumptive membrane-binding surface is at the bottom of the molecule.

### SI Appendix, Video S2

Hypothetical model for the conformational changes Sec7 undergoes when switching from the autoinhibited to active states. A model of the inactive conformation containing all residues of *T. terrestris* Sec7 was generated by SWISS-MODEL (10) templated with the cryoEM experimental model, and then a morphing transition was created using the AlphaFold predicted model of *T. terrestris* Sec7 in ChimeraX (11). The first few frames of this morph are looped in the beginning to highlight the presumed spontaneous, but infrequent, dissociation of the GEF domain from the HDS2 domain. Once Sec7 adopts the active conformation, it can be stabilized by binding to the activated forms of regulatory GTPases such as Arl1 (depicted in blue), allowing activation of the Arf1 GTPase substrate (depicted in red).

## Supplementary Methods

### Cloning of expression and purification constructs

All fragments used for cloning (insert and vector) were generated by PCR (Q5 polymerase, NEB Cat. No. E0555). When the template DNA carried the same selection marker as the final construct, it was linearized by a restriction enzyme that cleaved outside the desired amplicon prior to PCR to limit background transformants, and assembled by Gibson assembly (NEB Cat. No. E2621). *S. cerevisiae* Sec7 constructs used for complementation tests (plasmid shuffling assay) were cloned into the pRS415 vector with the Sec7 promoter and terminator. All yeast constructs are full-length and N-terminally GFP tagged, and otherwise only modified as indicated.

Purification constructs were cloned into the pPICZB vector. The full-length *T. terrestris* construct used for cryoEM was cloned from a plasmid containing *T. terrestris* Sec7 cDNA our group generated in a previous study (12), subcloned into pPICZB with 3C protease-cleavable 6xHis tags on the N and C termini. The full-length *S. cerevisiae* constructs were cloned into the same vector, but with an N-terminal 12xHis-GFP-Hrv3 tag. All constructs were verified by sequencing.

### Strain modification

Standard genetic techniques were used to generate yeast strains (SI Appendix, Table S2). To generate the *sec7*Δ*age2*Δ strain, BY4741 *sec7*Δ::KanMX (CFY409) was transformed with a pFa6 plasmid-templated (13) *age2Δ::HIS3* cassette using standard LiOAc transformation. Cells were grown to an OD of 0.6 in YPD, washed with water twice by centrifugation, then resuspended and incubated at RT in 100 mM Lithium acetate, 1 mM EDTA, 10 mM Tris pH 8.0 for 15 minutes (cells were concentrated ∼50 fold). 50 ul of this cell mixture was then added to 250 ul of transformation mix (30.5% PEG 3350, 100 mM LiOAc, 0.226 mg/ml salmon sperm DNA, 0.5-1 ug DNA) and incubated for 30 minutes with gentle agitation (rotating). For genomic integrations, 2 mM DTT was included in the transformation mix. The cells/transformation mixture was then heat shocked at 42 °C for 15 min. For drug selection, cells were washed and recovered for 3 hours in YPD before plating. For auxotrophic selection, cells were washed once in YNB with 5% YPD supplemented and plated immediately. *Pichia pastoris* expression strains were generated by transforming the KM71H strain using electroporation as described previously (14) after linearizing the pPICZB-Sec7 cassette with PmeI.

### Purification of *T. terrestris* Sec7 for cryoEM

After transformation, 5 colonies of CFY4970 were patched together on a fresh Zeocin plate, and cultured with autoinduction media as described(15). Cells were collected by centrifugation after 48hrs, and yeast cell paste was flash frozen in liquid nitrogen in small aliquots. Cell paste was then lysed in a Spex cryogenic mill (6875D) for 15 cycles, 15 cps, 2 min rest between cycles, and powder was stored at -80 until further processing. 20 g of lysate powder was thawed rapidly by adding 18 ml room temperature thaw buffer (55 mM Hepes, pH 7.4, 495 mM KOAc, 11% glycerol, 44 mM Imidazole, 1.1 mM DTT, 1.1 mM AEBSF, and 1.1 x Roche cOmplete Protease Inhibitor Cocktail) and sonicated with a macro-tip sonicator (100% power, 1s on 1s off) until mostly thawed (∼2 min). Then, 2 ml of 10% CHAPS was added (final concentration of 0.5%) and further sonicated (20s) to solubilize membranes. This lysate was clarified by centrifugation (20 min, 10,000 rpm in AV10 rotor, and then 30 min 20,000 rpm in SS- 34 rotor), and loaded onto a pre-equilibrated HisTrap column (Cytiva) with a syringe. Wash and elution was performed using an AKTA Pure with two buffers: Buffer A (50 mM Hepes, pH 7.4, 450 mM NaCl, 5% glycerol, 40 mM Imidazole, 1 mM DTT, and 0.1% CHAPS) and Buffer B (50 mM Hepes, pH 7.4, 300 mM NaCl, 10% glycerol, 500 mM Imidazole, 1 mM DTT, and 0.1% CHAPS). The column was washed with 25 CVs of 4% B, followed by a gradient to 35% B over 10 CVs to elute. Elution fractions were checked by SDS-PAGE, pooled, concentrated, flash frozen in liquid nitrogen, and stored at -80°C for later use.

### Purification of *S. cerevisiae* proteins for biochemical analysis

*S. cerevisiae* Sec7 constructs were grown, induced, and harvested as described above. Batch affinity purification was performed with resin that had been fragmented by sonication (see below). The clarified lysate was incubated with the resin for 2 hours at 4 °C with rotation. After binding the resin was washed five times with 10 ml wash buffer (50 mM Hepes, pH 7.4, 450 mM NaCl, 5% glycerol, 40 mM Imidazole, 1 mM DTT, and 0.1% CHAPS), and transferred to a fresh tube after the third wash. Sec7 was eluted from the NiNTA resin by 3C protease cleavage overnight at 4 °C. This elution was further purified by size exclusion chromatography using a Sepharose 6 increase 10/300 (SEC buffer: 25 mM Hepes, pH 7.4, 250 mM NaCl, 5% glycerol, 1 mM DTT). SEC fractions were analyzed by SDS-PAGE, pooled, concentrated, flash frozen in liquid nitrogen, and stored at -80 for later use. ΔN17-Arf1, myristoylated Arf1, and prenylated-Ypt31/GDI complex were purified as previously described(1, 8, 16).

### Preparation of fragmented affinity resin

In order to increase the surface area of the Ni-NTA resin, the resin was resuspended in water to make a 20% slurry, then sonicated at 80% power for three minutes with a macro-tip, (cycles: 20s on / 10s off). Following sonication we equilibrated the fragmented resin 5x with buffer at a lower speed (1000 rpm) to remove resin fines.

### CryoEM data processing

Movies were motion-corrected and dose-weighted using MotionCor2 (17), and micrograph defocus values were estimated using GCTF(18). Micrographs were manually inspected and culled to 3,401 usable micrographs, which were imported into CryoSPARC for particle picking (19). Defocus values were estimated with patch-CTF estimation in CryoSPARC and ‘blob picker’ was used to pick an initial set of particles. 2D-classification was used to generate templates for template picking. An initial set of 938,642 particles was used for Ab-initio model generation and 3D-classification was performed iteratively using heterogeneous 3D refinement to generate a final set of 296,177 particles.

These particles were re-extracted in RELION 3.1(20) and assigned GCTF-estimated per-particle defocus values. Iterative rounds of CTF refinement and Bayesian polishing improved the resolution from 7.14 Å to 5.3 Å. 3D classification was attempted, but no improvement was attainable for the dimer reconstruction. Focused refinement on a monomer(21) was performed using a mask including a single full monomer and a small portion of the other (corresponding approximately to the HDS4 domain) for particle subtraction after symmetry expansion. A second monomer mask containing only a single monomer was used for refinement. After several iterations of CTF refinement, fixed angle 3D classification isolated 196,888 symmetry expanded particles which produced a 3.7 Å (0.143 FSC) reconstruction after iterative CTF refinement. The published crystal structure of the DCB-HUS domain and AlphaFold prediction(12, 22) were used for guidance with *de novo* building in the few regions with poor side chain density. A monomeric atomic model was refined into the monomer map using Real Space Refine(23) in Phenix(24). A model for the dimer was then generated and refined into a composite dimer map produced from the monomer map and the consensus dimer map using Phenix Combine Maps, and validated in Phenix(24–26). See Tables 1 and 2 and SI Appendix, Figures S1 and S3.

For cryoDRGN analysis, we used TOPAZ to increase the likelihood of rare particles(27) (this did not improve resolution of the monomer). Starting with 1,240,146 topaz picked particles heterogeneous 3D classification in CryoSPARC generated a final stack of 280,528 particles that were then used to generate a dimer map with C2 symmetry imposed (6.3 A), followed by a focused monomer refinement without subtraction before cryoDRGN training and analysis (SI Appendix, Figure S2) (2).

### Fluorescence microscopy

Cells were grown overnight at 30 °C in liquid selection media (-Leu) to an OD of 0.6. Cells were allowed to settle on a coverslip dish (MatTek) for 10 min, and washed with fresh media. Imaging for SI Appendix, Figure S4 was done using a CSU-X spinning-disk confocal system (Intelligent Imaging Innovations) with a DMI6000 B microscope (Leica), 10031.46 NA oil immersion objective, and a QuantME EMCCD camera (Photometrics). Imaging was done using a DeltaVision Elite system equipped with an Olympus IX-71 inverted microscope, a DV Elite complementary metal-oxide semiconductor camera, a ×100/1.4 NA oil objective, and a DV Light SSI 7 Color illumination system with Live Cell Speed Option with DV Elite filter sets. Exposure and laser power were adjusted according to intensity, and were kept the same for all specimens being compared in an experiment. The brightness/intensity was equivalently adjusted across all images in an experiment using ImageJ.

### Liposome preparation

Synthetic TGN liposomes were prepared as described previously(16). In brief – lipid stocks in chloroform were combined in a pear-shaped flask to produce a lipid mixture mimicking that of the yeast TGN(28) (SI Appendix, Table S3). Chloroform was evaporated slowly in a rotary evaporator heated to ∼37 °C, then rehydrated in HK buffer (20mM HEPES pH 7.5, 150 mM KOAc) at 37 °C overnight. After gentle resuspension, the mixture was extruded through 100 nm filters 21 times and stored at 4 °C for no more than 1 month.

### Liposome flotation membrane binding assay

Liposome flotation was performed as described(29). Briefly – 100 nm liposomes were loaded with Arf1 by EDTA exchange of GMPPNP for a final concentration of 250 µM lipid and the indicated Arf1 concentration. Sec7 constructs were added to a final concentration of 550 nM, and incubated at room temperature for 1 hour. Then 2.5 M sucrose in HK (20mM HEPES pH 7.5, 150 mM KOAc) was added to 1 M final concentration, and 80 µl was transferred to a polycarbonate tube. Then a 100 ul layer of 0.75 M Sucrose was added, followed by 20 µl of HK. This was centrifuged at 20 oC in a TLA100 rotor for 30 minutes, and the top layer collected for SDS-PAGE analysis (12% acrylamide gel).

### In vitro Arf activation (GEF) assay

GEF activity was determined by measuring the native Tryptophan fluorescence of Arf1, as described previously(16). Briefly, to a final volume of 150 µl in HKM (20mM HEPES pH 7.5, 150 mM KOAc, 2 mM MgCl_2_, 1 mM DTT), 200 µM 100 nm TGN liposomes, Sec7 construct (with concentration as detailed below), 1 µM myristoylated Arf1, and 200 µM GTP were added to a quartz cuvette. For Figures 2 and 4, GEF was added to a final concentration of 100 nM. For Figure 6b, 20 nM GEF was added. For Figure 6a, 60 nM GEF was added. For Figure 5g and SI Appendix, Figure S6c, 200 nM GEF was added. Tryptophan fluorescence (297.5 nm excitation, 340 nm emission) was measured using a fluorometer, and between additions the reaction was well mixed and the fluorescence was allowed to stabilize. The order listed is the order components were added, except for reactions with preloaded Ypt31. For these reactions, 500 nM prenylated Ypt31 was loaded onto liposomes by EDTA exchange of GMPPNP at 30 °C for 30 minutes. The presence of GMPPNP in the liposome mixture dictated that Arf1 was added last. Curves were fitted in GraphPad PRISM 10 using a nonlinear regression (one phase association) accounting for drift if apparent in the data to extrapolate the rate constant, which was then divided by the GEF concentration to calculate the exchange rate.

## Notes

### Competing Interest Statement

The authors have declared no competing interest.

